# Deciphering the dynamical origin of the Adipogenic commitment during stress-induced disruption of the circadian rhythm

**DOI:** 10.64898/2026.09.18.752812

**Authors:** Debjit Das, Kajal Charan, Sandip Kar

## Abstract

Obesity is a major health concern worldwide and is caused by an excessive fat accumulation in the body due to adipogenesis. Recent studies suggest that excessive stress or disruption of circadian rhythms can enhance adipogenic commitment and lead to obesity. Moreover, this adipogenic commitment occurs at a specific time of day, highlighting a correlation with the circadian rhythm. However, the dynamical nature of this kind of stress-induced adipogenic differentiation remains an unresolved puzzle. Herein, we proposed a mathematical model centred on the key transcription factor PPARγ, associated with adipogenic differentiation, and demonstrated that adipogenic commitment is a bistable, irreversible differentiation process in which slow positive feedback regulation of PPARγ primarily controls the differentiation commitment threshold. The model reproduced the experimental observations regarding the extent of adipogenesis induced by constant and pulsatile stress input signals and explained the correlation between adipogenic differentiation time and circadian phase. It further predicts how to control the adipogenic commitment even under stressful conditions. Overall, our study elucidates how stress signalling dynamically governs adipogenesis.

## Introduction

Adipogenesis is a fundamental biological process through which preadipocytes (precursor fat cells) differentiate into mature adipocytes^1^. This differentiation process plays a pivotal role in maintaining energy and metabolic balance, thereby contributing to overall physiological health^2,3^. Every year, around 10% of fat cells normally renew^4^, and dysregulation of this process may lead to various metabolic disorders, primarily obesity, a growing health problem worldwide^5^. Recent studies suggest that stress from irregular or disrupted circadian rhythms can cause obesity^6,7^ by stimulating adipogenesis. This is caused by the disrupted regulation of a key hormone, glucocorticoid, which regulates several metabolic activities and oscillates in a circadian rhythm-dependent manner^8^. However, a disrupted circadian rhythm or chronic stress can flatten this oscillation^9^. Recent studies on the 3T3-L1 cell line show that the flattening of glucocorticoids, achieved by constant DMI (Synthetic glucocorticoid mimetic agent) input, can drastically increase adipocyte differentiation compared to oscillatory DMI input^9^. Intriguingly, studies further indicate that adipocytes differentiate within a specific time window during the day^10^, suggesting that adipocyte differentiation depends on a specific circadian phase. How distinct molecular regulators of adipogenesis and circadian rhythm orchestrate adipocyte differentiation, and how they cross-talk to optimally control adipocyte generation, remains poorly understood. Unravelling the mechanisms underlying the effects of disrupted sleep or stress on adipocyte differentiation will provide critical insights into the therapeutic control of obesity.

At the cellular level, adipogenesis is regulated by the key transcription factor PPARγ, whose activity is controlled by several molecular regulators through positive feedback interactions with varying time scales^11^. Experimental studies show that several adipogenic stimuli (rosiglitazone, pioglitazone, lobeglitazone, etc.) increase PPARγ expression^12^. As stimulus strength increases, higher levels of PPARγ promote adipocyte differentiation. Interestingly, at intermediate stimulus levels, both adipocytes and pre-adipocytes can coexist, indicating a bistable nature of the differentiation mechanism^1^. Recently, Zhang et al. have shown that adipocyte differentiation occurs during a specific phase of the circadian cycle, as determined by the dynamics of the circadian reporter Rev and the PPARγ threshold for differentiation^10^. The above-mentioned molecular regulations indicate that key adipogenic regulators interact intricately with circadian regulators to govern the dynamic organisation of adipogenesis.

Mathematical and computational modelling is one of the most promising tools for understanding complex dynamical processes. Consequently, several mathematical models have been developed to describe the dynamics of PPARγ during preadipocyte differentiation. Park et al. (2012) proposed a data-driven, quantitative model of adipogenesis^1^. Their work demonstrated that PPARγ levels can be bimodal, driven by three consecutive positive feedback loops among PPARγ, CEBPA, and CEBPB. Ahrends et al. further analysed the complex feedback framework that controls adipogenesis^11^. They highlighted how strong feedback interactions, along with cell-to-cell variability, shape cell fate decisions during preadipocyte differentiation. Recently, Zhang et al. (2022) incorporated a combination of fast and slow regulatory positive feedback components, namely CEBPA and FABP4, to capture the temporal evolution of PPARγ expression^10^. Although these studies provide valuable mechanistic insights into adipogenesis, most of these models were built to address specific experimental observations related to preadipocyte differentiation. We still require a simple yet general model of preadipocyte differentiation that provides a holistic understanding of adipogenesis by substantiating a wide range of experimental observations. Here, we endeavoured to produce an adipogenesis model that can systematically (i) explain how preadipocyte differentiation behaves under different stimulus conditions (such as stimulus pulses of varying duration and strength), (ii) capture its bistable nature, (iii) exhibit its overall statistical trends in adipogenesis under diverse stimulation conditions, and (iv) demonstrate the circadian gating behaviour.

In this study, we develop a minimalistic mathematical model centred on PPARγ regulation to capture the stimulus-dependent nature of adipogenesis. Our model captures the dose- and duration-dependent effects of the stimulus on adipogenesis, thereby indirectly mimicking both regular and disrupted circadian rhythms. Additionally, by incorporating circadian regulators and their dynamics into the model, we have explained the experimentally observed circadian gating phenomenon. Further, we have systematically analysed the roles of different interactions in this process of preadipocyte differentiation. Overall, our model provides a mechanistic understanding of adipogenesis, which can be used to elucidate the origins of obesity during circadian disruption.

## Model

The central regulator of adipogenesis is PPARγ, and its expression levels often indicate preadipocyte or adipocyte cellular states (**Fig. 1A**). PPARγ receives positive regulatory inputs from multiple proteins through either double-positive or double-negative feedback interactions^11^. Studies have shown that a combination of fast- and slow-regulating proteins (CEBPA and FABP4) is crucial for PPARγ dynamics^9^. The fast regulator (CEBPA) helps PPARγ maintain its rapid dynamics, while the slow regulator (FABP4) filters out oscillatory signals, thereby establishing a cell-intrinsic regulatory system for cell fate determination. PPARγ exhibits independent positive feedback regulation with both CEBPA and FABP4, reinforcing its own activation through each pathway^10^. Herein, we have built our overall model in 3 different stages. In our model, we have adapted the core network from the model proposed by Zhang et al. (2022), which includes both the positive feedback regulations of PPARγ (**Fig. 1B**). This activation of PPARγ involves an explicit dependence on the stimulus, as observed in the differentiation trends^9^. The stimulus in the PPARγ network corresponds to circadian-controlled glucocorticoid (GC) signalling. The glucocorticoid (DMI) input serves as the indirect activator of PPARγ (**Fig. 1B**). To better understand how direct PPARγ activation influences this process, we substitute the glucocorticoid input in the model with rosiglitazone, while maintaining the same underlying feedback network (**Fig. 1C**). Rosiglitazone directly binds with PPARγ and increases its activity^9^. In addition, it has been reported to increase PPARγ expression at the transcriptional level^13^. To capture these effects, we introduce an interaction into the model whereby rosiglitazone activates transcription of PPARγ mRNA (**Fig. 1C**).

**Fig 1.**
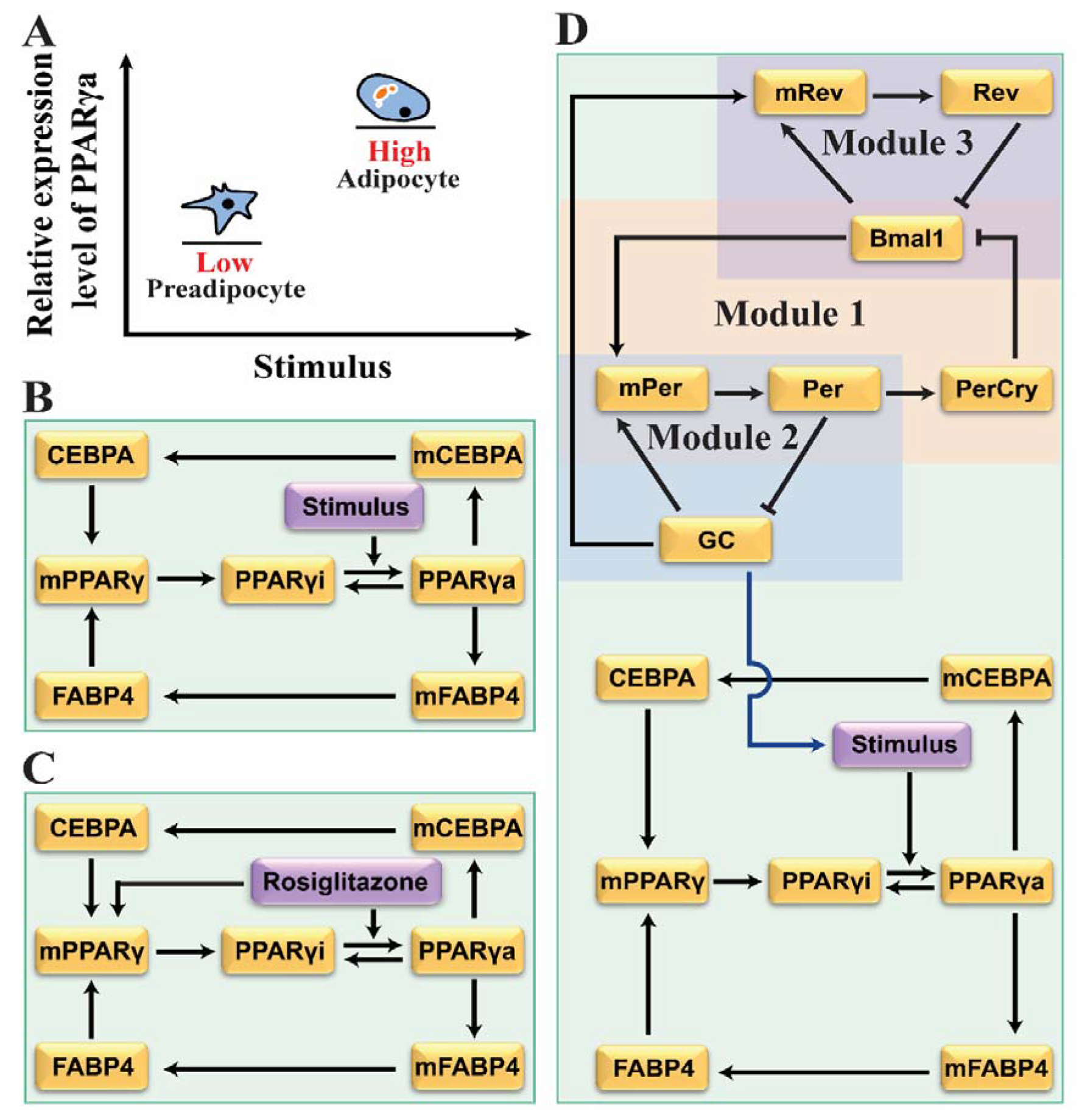
Models of stimulus-dependent Adipogenesis revolving around the central regulator PPARγ. **(A)** A schematic representation of the expression levels of PPARγ in preadipocyte and adipocyte cell states. **(B)** Molecular interaction network governing the stimulus (GC)-dependent PPARγ regulation. **(C)** Molecular interaction network governing the Rosiglitazone-dependent PPARγ regulation. **(D)** Proposed molecular interaction network model for the coupled circadian and adipocyte differentiation module. In all the diagrams, solid black arrows and hammer-headed lines indicate the biochemical activation and inhibition, respectively.

Further, to understand the interplay between circadian and adipogenesis, we have also included the circadian regulatory network (**Fig. 1D**), which can be divided into three modules. **Module 1** in **Fig. 1D** represents the core circadian oscillator, adapted from the models of Albert Goldbeter (1995)^14^ and Tyson et al. (1999)^15^. In this module, the Bmal1/Clock complex activates Per transcription, leading to the synthesis of the Per protein. Per subsequently forms a complex with Cry, and the PerCry complex then inhibits the Bmal1/Clock complex, thereby generating a delayed negative feedback loop that produces sustained circadian oscillations^16^. **Module 2** in **Fig. 1D** includes glucocorticoid (GC) signalling and its coupling to the circadian oscillator. In this module, GC enhances Per transcription, providing an additional regulatory input to the core clock^17^. The Per protein, in return, inhibits GC activity, forming a negative feedback loop between GC signalling and the circadian system^18^. This GC signal, generated by **Module 2**, serves as the upstream driver of stimulus in the adipogenic network. **Module 3** in **Fig. 1D** represents the Bmal1/Clock-Rev regulatory loop and how it gets modulated by GC signalling. In this module, the Bmal1/Clock complex activates Rev transcription, while the Rev protein inhibits the Bmal1/Clock complex, forming another negative feedback loop within the circadian system^19^. Rev functions as a circadian reporter whose expression remains unchanged between preadipocyte and adipocyte states^20^.

Rev expression is widely used as a readout of circadian phase during adipogenesis^6,21^. To mechanistically probe circadian gating within the model, we hypothesise that glucocorticoid signalling enhances Rev transcription, thereby coupling GC input to the core circadian machinery. This assumption allows glucocorticoid cues to modulate circadianphase-dependent repression and activation dynamics, thereby influencing the temporal window of adipogenic commitment. We have translated all the molecular interactions shown in these networks into deterministic, ordinary differential equation (ODE)-based mathematical models (**Table S1-3**). The related variables and parameters are explained in the **Table S4**.

## Results and Discussions

### Model captures bistability in PPARγ expression

To begin with, we have performed a systematic, deterministic bifurcation analysis to understand how PPARγ expression changes in response to a stimulus (**Fig. 2A**). In this context, the stimulus is considered to activate PPARγ (**Fig. 1B**), which is supported by experimental studies^1,9^. **Fig. 2A** depicts an irreversible bistable switch in PPARγ steady-state levels as a function of stimulus, indicating the presence of two distinct phenotypes in preadipocytes and adipocytes. Starting with a low PPARγ value (a preadipocyte), as the stimulus increases, PPARγ levels rise. At the saddle-node point (**S**_**N**_**1**), the system undergoes a transition to the upper steady state, corresponding to the differentiated adipocyte state. This is consistent with experimental studies showing that PPARγ levels are directly associated with adipocyte fate: low PPARγ levels are found in undifferentiated preadipocytes, whereas high PPARγ levels are found in differentiated adipocytes^1^. Importantly, some signals (such as DMI, rosiglitazone, pioglitazone, and lobeglitazone) can drive this process^22–25^.

**Fig 2.**
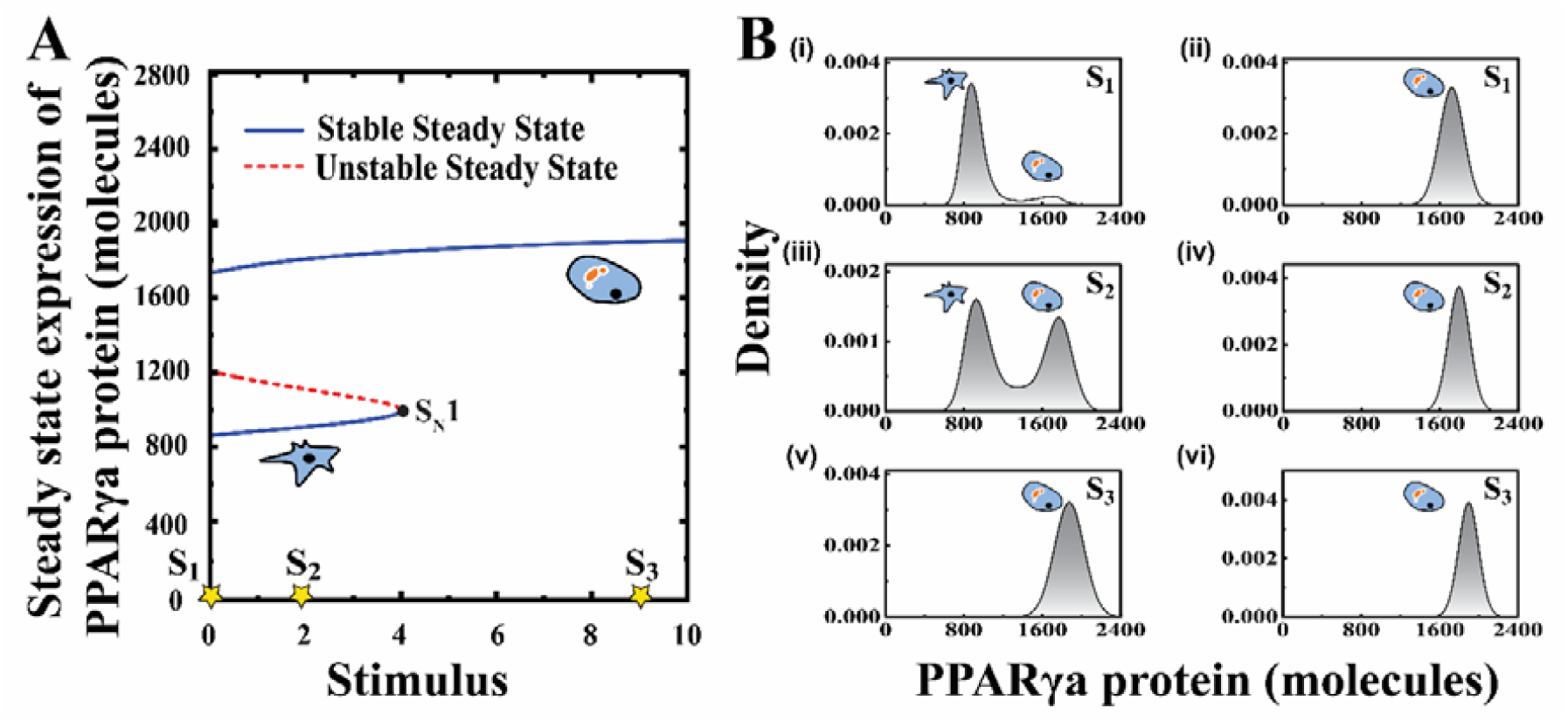
Adipogenesis happens via an irreversible bistable switch of PPARγ expression level as a function of Stimulus. **(A)** Deterministic bifurcation of PPARγ steady state as a function of stimulus. **(B)** Distribution of PPARγ protein at different stimulus values (**S**_**1**_=0, **S**_**2**_=1.9, **S**_**3**_=9) (Stochastic simulation for n=4500 trajectories), starting from low initial condition (left panel, **(i), (iii)** and **(v)**) and from high initial condition (right panel, **(ii)**, and **(vi)**).

Further, the model predicts that once the system attains a higher steady-state level of PPARγ, a subsequent decrease in stimulus does not revert PPARγ to its low level (as the 2^nd^ critical point (**S**_**N**_**2**) lies in the negative domain of the stimulus). It implies that cells, once differentiated, remain in that state even after the stimulus is removed, demonstrating the irreversibility of differentiation. This means that adipogenesis is a terminal differentiation (**Fig. 2A**). However, under intermittent signal strength, the system can exist in either a preadipocyte or an adipocyte phenotype. Intriguingly, in the experiments, higher signal strength triggers differentiation, with only adipocytes observed, whereas at low signal levels both preadipocytes and adipocytes can coexist, demonstrating the bistable nature of the process^1^, as predicted by our simple model. Moreover, cells that differentiate into adipocytes maintain high PPARγ expression during the post-differentiation period, suggesting that adipogenesis may be an irreversible bistable process^1^. Next, to determine whether both steady states within the bistable window could be captured at the single-cell level, we converted our ODE-based mathematical model into a stochastic model using Gillespie’s stochastic simulation algorithm^26,27^. Stochastic simulation was performed for 4500 cells at various stimulus values (**Fig. 2B**). For cells starting from preadipocyte-like (low PPARγ levels), as stimulus levels increase, the fraction of cells that undergo adipogenesis increases, and at intermediate stimulus levels, both states coexist **(Fig. 2B, left panel)**. In contrast, simulation starting from an adipocyte-like state or from high PPARγ levels does not return to a low PPARγ level, indicating the irreversibility of the system **(Fig. 2B)**. Once adipocytes are formed, the reverse process of converting into pre-adipocytes is not feasible. The duration of 4 days is chosen to match typical experimental time frames, ensuring the relevance of the simulation results to experimental conditions^9^ **(Fig. 2B)**.

### FABP4 drives PPARγ after crossing the commitment threshold

Such differentiation dynamics, dependent on PPARγ expression level, have two characteristic features that are extremely crucial to understand. First, beyond which threshold level of PPARγ can we safely consider that a pre-adipocyte cell has converted into an adipocyte? Secondly, what factors optimally control the timing of differentiation? Zhang et al.^10^ used the Receiver Operating Characteristic (ROC) method to experimentally demonstrate that commitment of pre-adipocytes to adipocytes is facilitated by a threshold level of PPARγ. We have also performed ROC analysis (**Method**) and estimated a threshold for active PPARγ molecules **(Fig. 3A-B)** to numerically designate the pre-adipocyte-to-adipocyte transition. The transition from pre-adipocyte to adipocyte is facilitated by the switch-like activation of PPARγ, which is governed by positive feedback regulation from CEBPA and FABP4.

**Fig 3.**
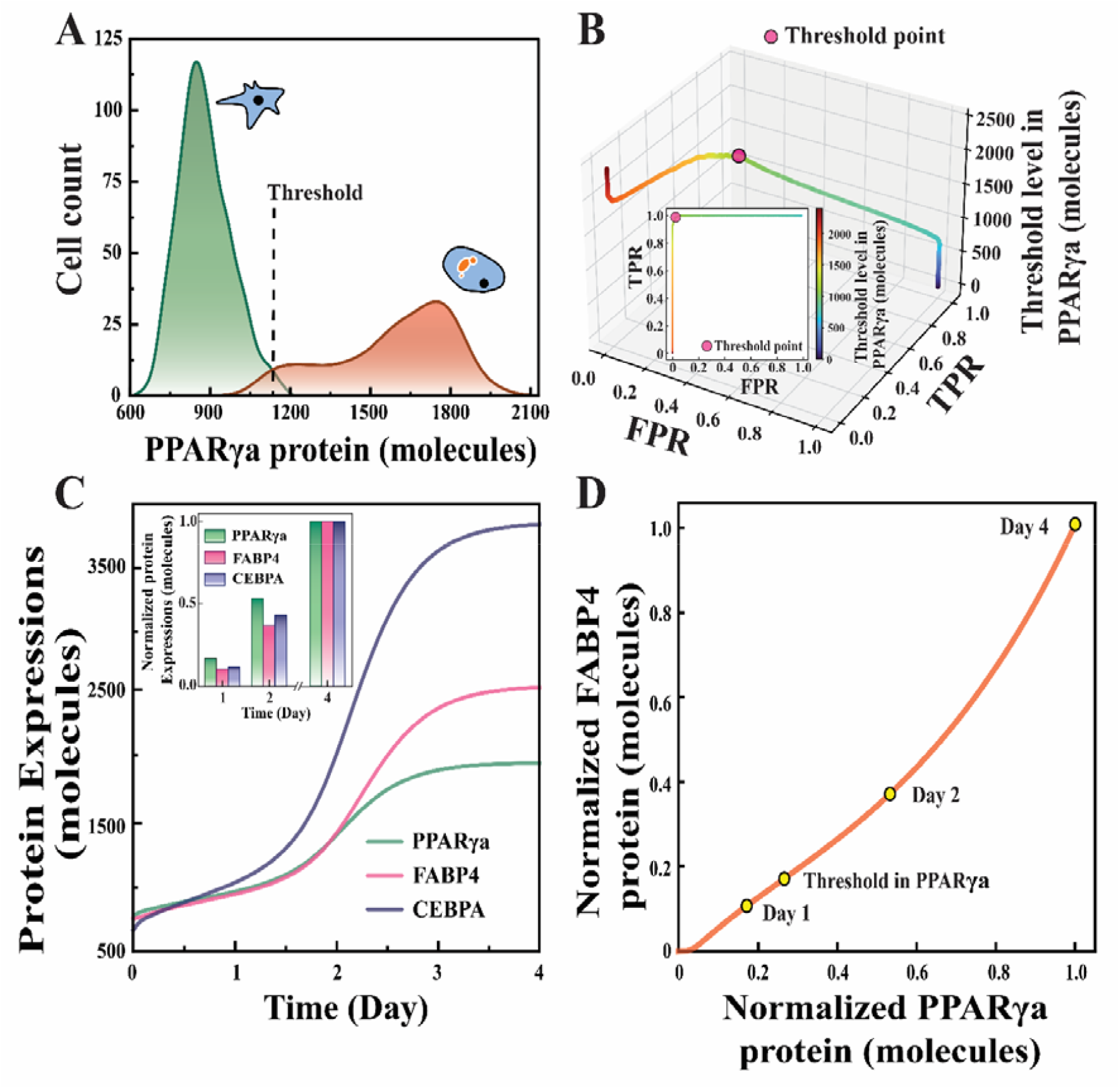
FABP4 governs the late rise of PPARγ after crossing the commitment threshold for adipogenesis. **(A)** Schematic representation of bimodality in PPARγ expression level simulated for stimulus value = 1.9. **(B)** ROC analysis plot between True positive rate (**TPR**) and False positive rate (**FPR**) with threshold levels of PPARγ represented in 3D (inset shows the same in 2D representation). **(C)** Protein expression levels of PPARγ, CEBPA and FABP4 for 1, 2 and 4 days under stimulus =25 condition (**inset** shows the normalised protein levels under the same stimulus condition). (D) Normalised levels of PPARγ and FABP4 plot for 4 days of simulation.

The nature of activation varies in terms of timescale: CEBPA is a fast regulator, whereas FABP4 is a slow regulator protein. Identifying the threshold level of active PPARγ numerically has allowed us to understand how these temporally distinct feedback loops shape the differentiation time course. Interestingly, between days 1 and 2, induction of CEBPA is much higher than that of FABP4, and levels of active PPARγ remain moderate (**Fig. 3C, inset**), suggesting that PPARγ activation precedes full FABP4 accumulation. Indeed, our simulation results reveal that the sharp increase in FABP4 expression temporally coincides with the PPARγ threshold-crossing event (**Fig. 3C**). Specifically, as FABP4 levels begin to rise steeply, PPARγ crosses its commitment threshold and irreversibly transits to the adipogenic state^9^. This temporal coupling indicates that while PPARγ shows rapid dynamic activity, effective and sustained threshold crossing depends on adequate accumulation of FABP4. These model outcomes are in line with the experimental studies, which show that the CEBPA possesses a fast turnover rate comparable to that of PPARγ and therefore supports rapid PPARγ dynamics through a fast positive feedback mechanism^28^, while FABP4 exhibits slow turnover kinetics and accumulates gradually over time, introducing a delayed regulatory component into the system^29^.

To better visualise the correlation between FABP4-induced PPARγ activation, normalised FABP4 is plotted against normalised PPARγ (**Fig. 3D**). During the early phase, the trajectory lies closer to the PPARγ axis, indicating that the system dynamics are initially dominated by PPARγ activation via CEBPA, which is later guided through FABP4 accumulation. Following the threshold-crossing event (**Fig. 3D**), the trajectory approaches the diagonal, reflecting synchronised late activation of both proteins. These results indicate that while CEBPA maintains rapid PPARγ dynamics, the delayed accumulation of FABP4 governs the effective threshold-crossing event. Thus, PPARγ commitment is not driven solely by its intrinsic fast feedback loop, but is critically guided by the slow regulation of FABP4.

### Sustained input of DMI significantly enhances the extent of adipogenesis

Next, we wanted to investigate whether or not our model could differentiate between oscillatory and sustained dynamical inputs to capture the circadian rhythm’s effect on adipocyte differentiation. Experiments have shown that DMI, a potent synthetic glucocorticoid, when given as an oscillatory input (12 hours on and 12 hours off), results in a low adipocyte count. However, sustained DMI input increases adipocyte count significantly^9^. We conducted a simulation protocol following the experimental schemes by introducing a pulse stimulus into our model. In these simulations, the total stimulus dose varies with pulse duration **(Fig. 4A)**.

**Fig 4.**
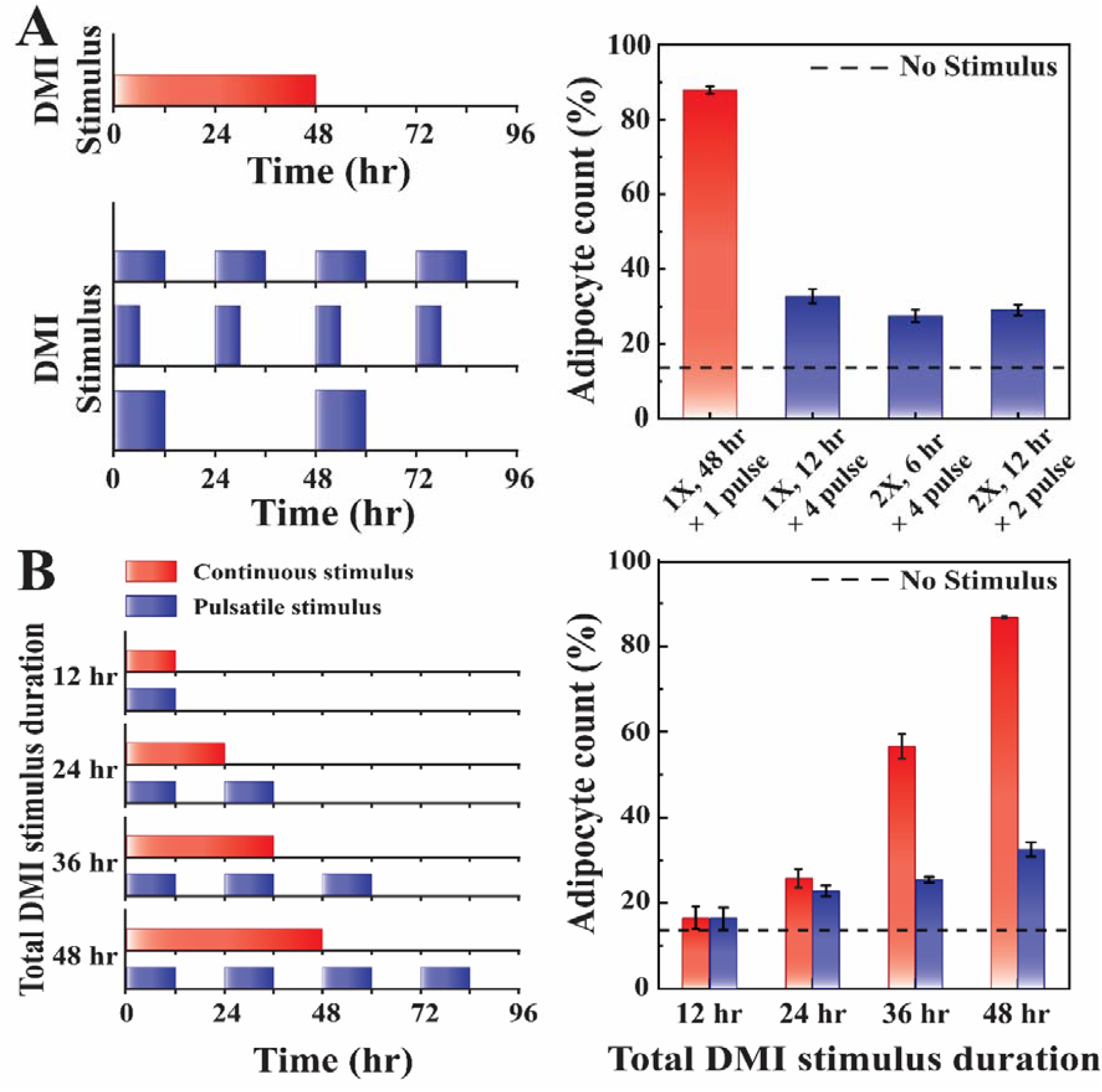
Sustained DMI input significantly enhances the extent of adipocyte differentiation in comparison to the pulsatile input. **(A) (i)** Protocol for numerically altering the DMI stimulus inputs (total constant DMI with varying dose strength and time interval).**(ii)** Percentage adipocyte count from the simulations under different DMI inputs (error bar indicates standard deviation).**(B) (i)** Protocol for numerically altering the DMI stimulus inputs (fixed total DMI for pulsatile and sustained stimulus). **(ii)** Percentage adipocyte count from the simulations under different DMI inputs (error bar indicates standard deviation).

We ran simulations for four sets with a fixed total stimulus dose. In the first case, where the stimulus is provided as a constant input **(Fig. 4A(i))**, a high (∼87%) adipocyte count is obtained **(Fig. 4A(ii), Red bar)**. In the other three cases **(Fig. 3A(i))**, the stimulus is delivered in pulses of varying duration and intensity while maintaining the total stimulus dose constant.

It is observed that the adipocyte count for all three different DMI pulsatile inputs **(Fig. 4A(i))** is ∼55% lower than the constant DMI input condition **(Fig. 4A(ii)**. Further, we have examined what happens when the pulse ON time is kept similar but with different patterns **(Fig. 4B (i))**. Our simulations demonstrate that over 24 hours, there is an insignificant difference in adipocyte count **(Fig. 4B (ii))**. However, for the 36 and 48-hour cases, a constant stimulus significantly alters adipocyte count compared to a pulsatile stimulus **(Fig. 4B (ii))**. These simulation results corroborate the experimentally observed trends under different DMI stimulus conditions^9^.

This suggests that differences in adipocyte differentiation are not solely driven by the total amount of stimulus exposure. Instead, the temporal behaviour of the stimulus input plays a crucial role in determining the extent of differentiation. Thus, our model clearly distinguishes the disparate nature of differentiation due to prolonged and oscillatory DMI signals. Continuous signals effectively drive the system towards differentiation by maintaining activation for a sufficient period. However, any pulsatile signal fails to provide sustained activation and is less effective for differentiation.

#### Direct activation by rosiglitazone triggers adipogenesis even with a short pulse stimulus

In the previous section, we saw that DMI input indirectly activates PPARγ via CEBPB^9^, and that sustained DMI input leads to greater adipogenesis than pulsatile DMI input. In a similar context, experimental studies have shown that when PPARγ is directly activated by rosiglitazone (**Fig. 1C**), adipocyte count increases with increasing stimulus pulse duration (**Fig. 5A**), and even with a 4-hour rosiglitazone pulse, there is a considerable increase in adipogenesis, unlike the DMI pulse input^9^. The differentiated cell fraction continued to increase with longer pulse durations and approached saturation at around 24 hours (**Fig. 5A**), at which nearly all cells had committed to the differentiated state^13^. This rapid, saturating response highlights the greater efficacy of rosiglitazone in directly activating PPARγ in driving adipogenesis compared with the slower, delayed response observed with indirect activation via DMI. To compare the difference between direct and indirect regulation effects on adipocyte count, we have also simulated with varying DMI pulse durations (**Fig. 5A**) similar to the rosiglitazone. We observe that after 4 days of DMI stimulation, the differentiation percentage remains constant for 12-16 hours, then increases progressively **(Fig. 5A)**, consistent with the experimental observation^9^.

**Fig 5.**
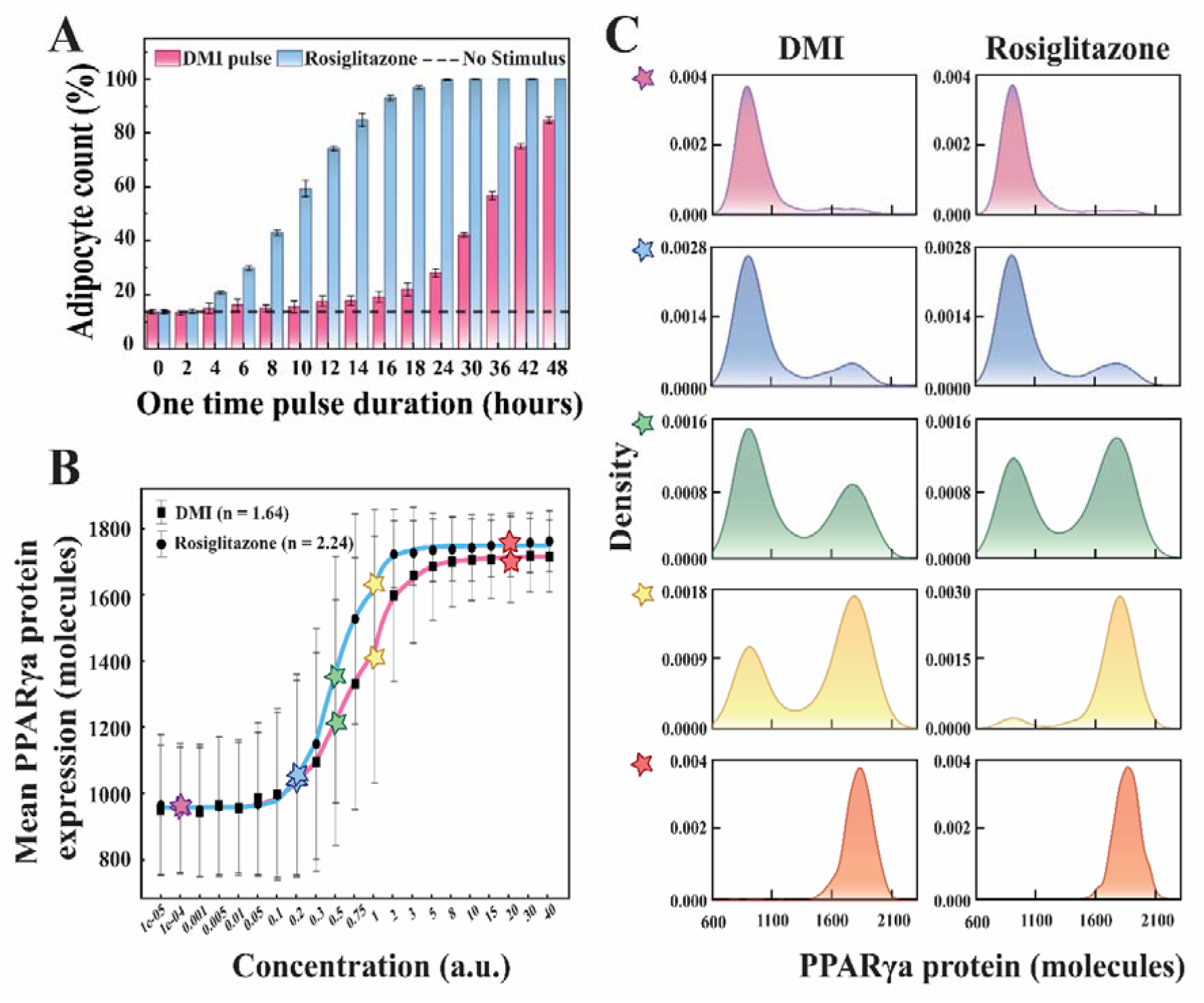
Rosiglitazone activates adipogenesis with greater cooperativity. **(A)** Percentage adipocyte count obtained for different one-time pulse durations (ranging from 0 to 48 hr) of DMI and Rosiglitazone. **(B)** Dose response of mean PPARγ protein expression as a function of Rosiglitazone and DMI concentrations. The blue and red solid lines denote the fitted lines with Hill functions (having Hill coefficient n=2.24 and n=1.64) for Rosiglitazone and DMI, respectively, while the black dots and squares represent the mean PPARγ expression levels from each simulation. **(C)** Distribution of PPARγ protein expression (∼1200 individual stochastic trajectories) across different rosiglitazone levels (as indicated by colored stars in **Fig. 5B**).

To understand how direct PPARγ activation by rosiglitazone differs from indirect DMI input, we used the model developed based on the network shown in **Fig. 1C**. To quantitatively characterize it, we have numerically obtained the dose-response of the mean terminal PPARγ expression as a function of rosiglitazone and DMI concentrations (which is varied over a broad range 10^-5^ to 40 a.u.), respectively **(Fig. 5B)**.

The simulated steady-state PPARγ response exhibits a sigmoidal dependence on rosiglitazone concentration **(Fig. 5B)**, which reconciles the experimental findings of Park et al. Fitting the corresponding dose-response curve with a Hill function via nonlinear regression results in a Hill coefficient of 2.24 **(Fig. 5B)**^1^. The Hill coefficient greater than unity indicates positive cooperativity in the effective activation of PPARγ by rosiglitazone. Our model simulation predicts that the steady-state of PPARγ will also vary sigmoidally as a function of DMI concentration, however, the dose-response curve can be fitted by a Hill function with a smaller Hill coefficient (1.64) **(Fig. 5B)**. This suggests that the rosiglitazone-mediated activation is amplified by the network-level regulatory interactions to a greater extent, which leads to a higher cooperative response in comparison to DMI-mediated activation. This can be further confirmed by observing the bimodal PPARγ expression pattern at moderate rosiglitazone and DMI doses **(green and yellow stars, Fig. 5C)**. Such an early rise of cells with higher PPARγ steady-state levels indicates that rosiglitazone can induce adipogenic commitment much earlier due to higher extent of cooperative feedback dynamics in comparison to DMI-mediated activation.

#### Glucocorticoid-mediated Rev transcription drives the circadian gating of adipogenesis

Thus far, the model has captured distinct signal-pattern dependencies that mimic circadian rhythm inputs in adipocyte differentiation. Under normal physiological conditions, adipose tissue undergoes dynamic turnover, with around 10% of fat cells renewing each year to maintain a steady fat content^4^. Intriguingly, this differentiation occurs within a specific time window during the day. Experimental studies using the OP9 cell line with the Rev (circadian reporter) and PPARγ proteins indicate that differentiation occurs at a specific phase of the circadian rhythm. This suggests that the adipocyte differentiation network intrinsically exhibits circadian gating, with commitment occurring preferentially during a specific phase of the circadian cycle^10^. This raises two important questions: (i) how does the circadian rhythm interact with the regulation of adipocyte differentiation, and (ii) what are the key molecular nodes involved? To address these questions, we extended our adipocyte differentiation network to incorporate key circadian interactions (**Fig. 1D**, explained in the model section).

In our simulation, the differentiation time of an individual cell was defined as the time at which its PPARγ protein level irreversibly crossed the commitment threshold. The corresponding phase of the Rev oscillation at this time point was then assigned as the differentiation phase of adipogenic commitment for the respective cell (**Fig. 6A**). This analysis is in line with a standard experimental protocol proposed by Zhang et al.^10^. Rev expression is intrinsically oscillatory and remains qualitatively unchanged between preadipocytes and adipocytes, making it a suitable phase marker^20^. Here, each peak of the Rev oscillation was assigned a phase of 0 or 2π, while each trough was assigned a phase of π.

**Fig 6.**
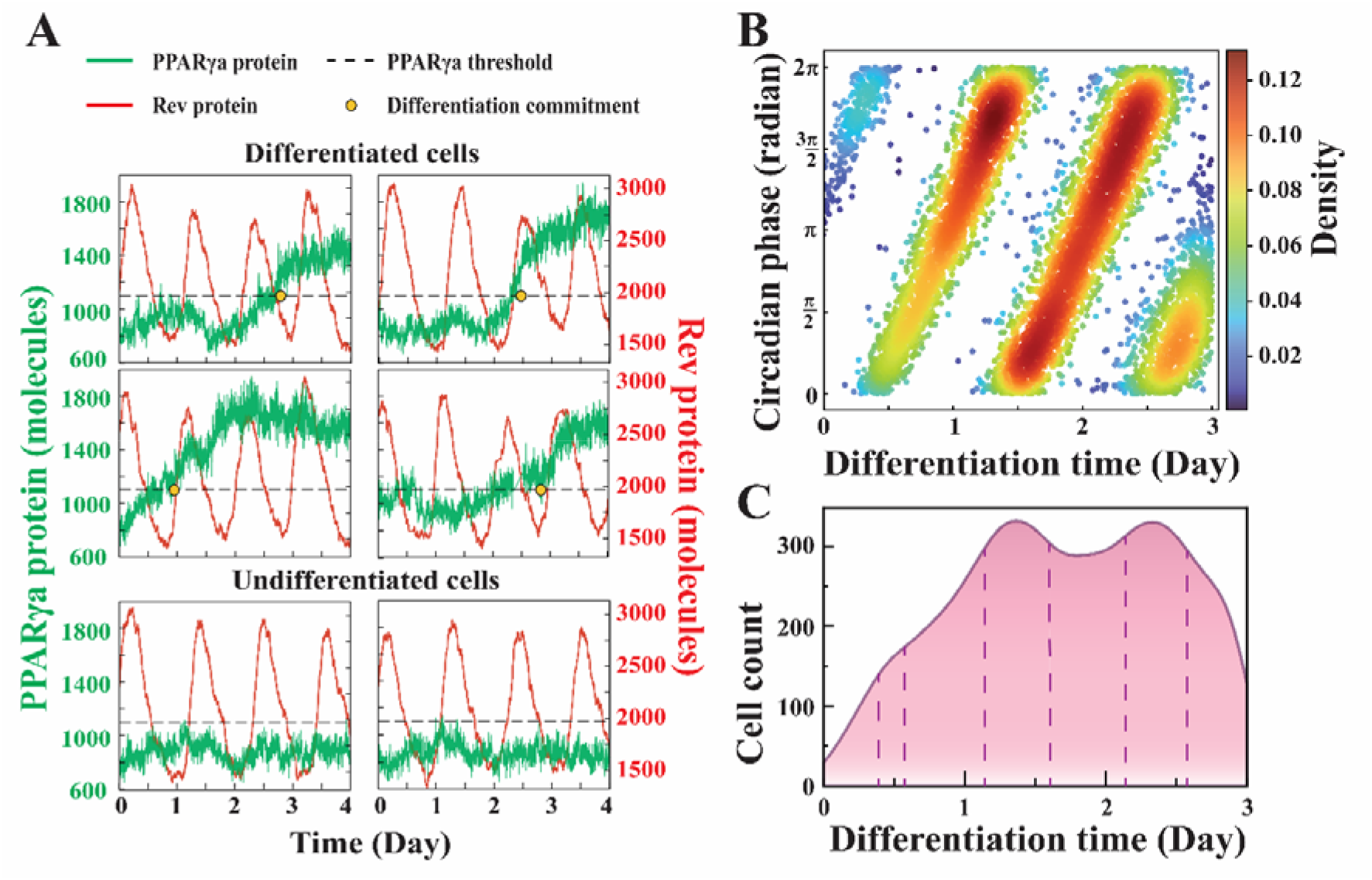
Model reconciles the circadian gating of Adipogenesis. **(A)** Representative time profiles of Rev and PPARγ levels for differentiated (top panel) and undifferentiated (bottom panel) cells from stochastic simulation.(B) Density plot for circadian phase and differentiation time for 7520 trajectories. **(C)** Cell count versus differentiation time-of-day plot for 7520 trajectories.

Consequently, the interval from 0 to π corresponds to the falling phase, whereas π to 2π represents the rising phase of the circadian cycle. Our numerical model simulations reveal that most adipogenic commitment events occur during the rising 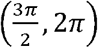 phase of the circadian cycle (**Fig. 6A**), which reconciles the experimental observations made by Zhang et al.^10^. Although there is a slight phase shift in the timing of commitment relative to the experimental observations, this provides the crucial mechanistic insight that intrinsic network dynamics bias adipogenic commitment toward a specific circadian phase.

Importantly, our model analysis unravels that this phase restriction emerges from the incorporation of glucocorticoid-driven Rev transcription. In the absence of this interaction, Rev oscillates according to its intrinsic circadian rhythm and remains phase-shifted relative to glucocorticoid input. As a result, differentiation commitment was found to form a uniform distribution with respect to the Rev phase. Upon introducing transcriptional enhancement of Rev by glucocorticoid, these two circadian components become phase-aligned, exhibiting coincident peaks and troughs. This synchronisation effectively entrains the Rev oscillator to the glucocorticoid signal, reducing phase lag and stabilising their temporal relationship (**Fig. 6A**). Most adipocytes differentiate during the rising phase of the Rev. Consequently, the density plot in **Fig. 6B** also shows a high density at this 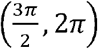 phase. Additionally, the proportion of differentiated cells was high during the rising phase. There appears to be heterogeneity in the timing of cell commitment after day one **(Fig. 6B)**, suggesting that both the specific circadian phase and the levels of intrinsic regulators are important. The specific circadian phase may trigger this process, but the delay in threshold crossing can result from the relative levels of other regulators. However, the timing of differentiation peaks at specific intervals and times of day, as shown in **Fig. 6C**.

#### Reducing pulse sensitivity can decrease the rate of adipogenesis even at a constant pulse

Irregular sleep patterns have become part of our daily lives, potentially flattening the rhythmic GC dynamics normally driven by the circadian cycle. It leads to a continuous GC signal that significantly increases the rate of adipogenic commitment. As a result, adipose tissue becomes enriched with adipocytes, and thus the individual becomes obese. Herein, we aim to identify ways to alter the differentiation outcome, even under a constant GC signal, by modulating biological interactions in our model. Specifically, we investigated how the underlying biological network that regulates adipogenesis distinguishes between continuous and oscillatory GC signals. significantly change the extent of differentiated cells in the case of pulsatile signals (4-6.5% relative decrease) (**Fig. 7A**).

**Fig 7.**
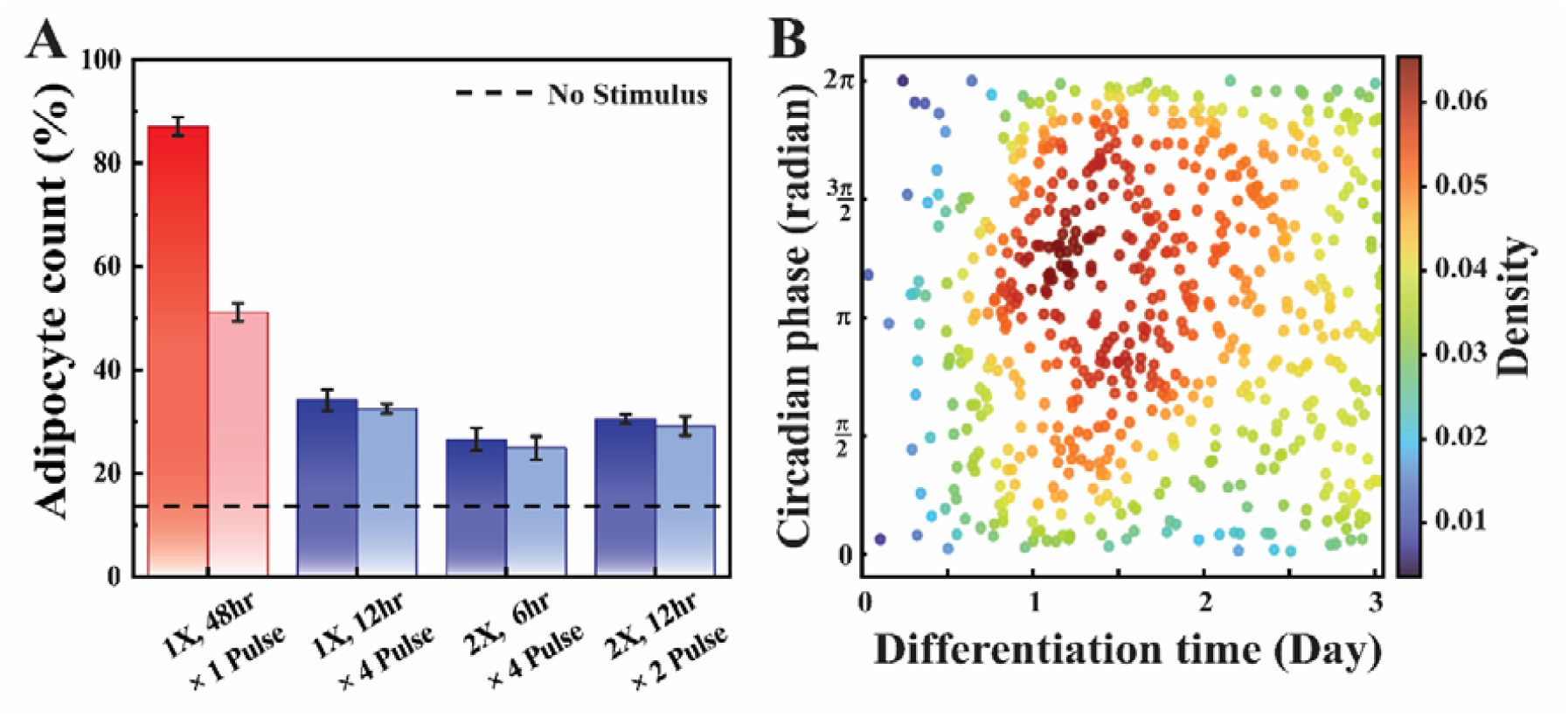
Model predicts ways to control Adipogenesis and its related consequences. **(A)** Reduced pulse sensitivity markedly decreases adipocyte count, whereas adipocyte formation remains unaffected under pulsatile stimulation. (B) Continuous stimulation abolishes the circadian phase-differentiation time correlation observed under pulsatile signalling.

Our model analysis reveals that reducing the feedback strength of either CEBPA or FABP4 results in a significant decrease in adipocyte count (**Fig. S1**). However, this effect is persistent across all signal patterns (**Fig. 4A**). Even for pulsatile signals, the extent of differentiation extends beyond the no-stimulus condition. Thus, by experimentally perturbing these feedback regulations, we cannot reduce the extent of adipogenesis specifically under constant GC signalling. However, by reducing the sensitivity to the pulse duration by up to ∼50%, the adipocyte count drops from 87% to ∼51% (44% relative decrease) for a 48-hour continuous signal (**Fig. 7A**). Such a reduction in sensitivity to pulse duration does not

#### Flattening of GC signalling diminishes circadian gating during Adipogenesis

An interesting feature of adipogenesis is circadian gating. We have previously shown that there is a strong correlation between circadian phase and differentiation timing, and that most adipogenic differentiation commitment occurs during the rising phase of each circadian cycle. Generally, GC restores its expression level at night, but irregular sleep or stress can disrupt this rhythm, leading to a prolonged GC signal. Considering this situation, we performed a similar numerical simulation to obtain the circadian phase vs. differentiation time plot (**Fig. 7B**). We observed that the maximum differentiation commitment still occurs during the rising phase of the first circadian cycle (**Fig. 7B**). However, the correlation between circadian phase and differentiation time is no longer present (**Fig. 7B**). Since the preadipocytes are now exposed to continuous GC signals, the rate of adipogenesis is so high that the cells underwent early differentiation (during the first circadian cycle since the constant signal is activated), that disrupts the correlation of circadian phase and the differentiation time leading to a random distribution.

## Conclusion

Every year, around 10% of fat cells differentiate via adipogenesis, ensuring the adipose tissue homeostasis^4^. The disruption in this tightly regulated process due to chronic stress or poor lifestyle can contribute to obesity and other metabolic diseases. The underlying molecular regulatory network for adipogenic commitment at the cellular level is primarily governed by the transcription factor PPARγ. In this article, we have performed a systematic modelling study (**Fig. 1**) and proposed that the phenotypic transition from preadipocyte to adipocyte is characterised by a bistable steady-state expression of PPARγ, in which preadipocytes express low levels of PPARγ and adipocytes express high levels of PPARγ (**Fig. 2**). Our model captures both bistability in PPARγ expression and irreversibility of preadipocyte to adipocyte transition (**Fig. 2**) and elucidates the mechanistic role of fast and slow positive feedback loops in PPARγ activation during differentiation (**Fig. 3**). The model analysis explains that early rise in PPARγ leads to the slow accumulation of FABP4, and once PPARγ crosses the threshold of differentiation, the FABP4-positive feedback helps maintain its high level. The relative timescale of these regulatory interactions is critical to determining how the system responds to oscillatory and sustained stimulus inputs (**Fig. 3**).

Consistent with experimental observations^9^, our model distinguishes between sustained and oscillatory glucocorticoid inputs (**Fig. 4**). Continuous stimulation efficiently drives differentiation, whereas pulsatile signals, even with an equivalent total dose, are markedly less effective. This provides a mechanistic explanation for the experimentally observed dependence of adipogenesis on the temporal structure of the stimulus rather than on stimulus magnitude alone (**Fig. 4**). However, the direct PPARγ activator, Rosiglitazone, does not exhibit this filtering (**Fig. 5**) and instead shows a pulse-duration-dependent increase in adipocyte count^9^. Our simulations suggest that rosiglitazone’s higher positive cooperativity in the effective activation of PPARγ, compared with DMI (**Fig. 5B-C**), enables it to bypass upstream filtering and achieve rapid commitment even under short-duration pulses.

We have further extended the model by coupling it with the circadian module to find out the origin of the circadian gating phenomenon in the context of adipogenesis. Our model analysis reveals that circadian gating arises from glucocorticoid-driven transcriptional regulation of Rev, which synchronises hormonal input with the core circadian oscillator (**Fig. 6**). Entrainment of Rev biases differentiation toward a specific circadian window, providing a network-level mechanism for circadian gating. Our model shows that, despite being a heterogeneous process (**Fig. 6B**), it captures cells’ bias towards the rising phase of the rev, consistent with experimental observations^10^. Intriguingly, the model predicts that obesity (greater accumulation of adipocytes) under stressful or sleepless conditions can be avoided if the system remains less sensitive to the signal’s pulse duration (**Fig. 7A**). It further reveals that, under such conditions, circadian gating will be lost (**Fig. 7B**).

Overall, our modelling study has provided a mechanistic rationale about how adipogenic commitment depends on the strength and nature of signalling inputs. The model further explained how stress or disturbed sleep patterns govern obesity and how to restrict adipogenic differentiation even under stressful conditions. We believe that these insights will be useful for generating therapeutic strategies to tackle obesity-related problems in the future.

## Material and method

### Effective activation rate constant at different stimuli

We hypothesise that PPARγ activation can sense the duration of stimulus pulses, rather than responding solely to instantaneous signals. This assumption reflects the ability of the regulatory machinery to distinguish between transient and sustained inputs, which is particularly relevant to oscillatory glucocorticoid signalling^9^. In the model, the stimulus is represented as a periodic pulse of fixed strength and duration. Specifically, the stimulus S(t) is defined as

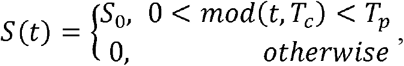

where S_0_ denotes the pulse strength, T_c_ is the total pulse duration (corresponding to one circadian period), and T_p_ is the pulse duration within each cycle. Moreover, to account for stimulus-dependent regulation within the network, we replace the basal activation rate constant (k_act_) of PPARγ with an effective, pulse-duration-dependent activation rate constant that explicitly incorporates the stimulus effect.

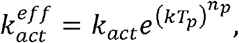

Where k is a scaling parameter that sets the sensitivity to pulse duration and n_p_ controls the degree of non-linearity.

**Deterministic Simulation**

Our deterministic model includes 14 ODEs that describe the dynamical behaviour of mRNAs and proteins of the interaction network motif. Bifurcation analysis is conducted using a freely available software XPPAUT (https://sites.pitt.edu/∼phase/bard/bardware/xpp/xpp.html).

### Stochastic Simulation

Cells behave differently under identical physiological conditions. However, deterministic simulations describe the average behaviour of the system; they overlook molecular fluctuations that arise from low copy number or probabilistic events. Therefore, we extend our analysis by implementing Gillespie’s stochastic simulation algorithm^26,27^ to capture the intrinsic randomness associated with adipogenesis. This algorithm is based on a chemical master equation formalism, where each biochemical reaction occurs with a probability determined by its propensity. In our stochastic simulation, we have converted the scaled units to molecular number by introducing a scaling factor. Each numerical simulation picks up a different random number seed which includes cell-to-cell heterogeneity and allows us to capture the full spectrum of stochastic behaviour. To examine how the stimulus timing influences adipogenic differentiation, simulations were performed from day -1 to day 4, with stimulation initiated at day 0 in each case. This setup allows the system to reach a comparable initial state before stimulation begins. Therefore, differences in differentiation fate arises from stimulus patterns rather than from initial conditions.

### Threshold Determination: ROC Analysis

The crucial step in adipogenesis process is finding the threshold level of the main regulator PPARγ. The most crucial aspect of this whole study was the determination of differentiation commitment to a cell. A fat cell may exist as either of the two phenotypes. Such binary decision problems have a threshold depending on which the fate is decided. To determine the threshold in PPARγ level we employed Receiver Operating Characteristic (ROC) curve-based approach, maintaining consistency with the experimental methodology^10^. This methodology is applicable when the dataset has two distributions with significant overlap (Fig 2A (i)). Depending on the threshold we may classify the possible outcomes of the process into four classes: (i) True Positive (TP): If the actual event is true and the model also predicts it as true, then it is a true positive case, (ii) True Negative (TN): If the actual event is false and the model predicts it as false, then it is a true negative case, (iii) False Positive (FP): If the actual event is false but the model predicts it as true, then it is a false positive case, (iv) False Negative (FN): If the actual event is true but the model predicts it as false, then it is a false negative case. Depending on the requirement of the model, we prioritize to reduce FP or FN so that the model can predict the true events correctly and report the false ones with confidence.

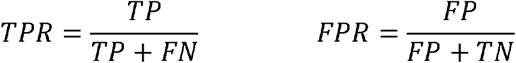

Now we determine two metrices for the model with the help of these four parameters: Sensitivity or True Positive Rate (TPR) of the model is the probability of identifying the true positive events correctly out of all actual positive events. Specificity of the model is the probability of identifying the true negative events out of all actual negative events. And False Positive Rate (FPR) is (1-Specificity). Now the ideal case scenario for a model is when the model maximizes TPR along with minimizing FPR, i.e., (0,1) point in TPR vs FPR plot. In reality, the coordinate in TPR vs FPR plot that lies the closest to this (0,1) point corresponds to the threshold of the system (Fig 2A (ii, iii)).

## Acknowledgments

Thanks are due to UGC for providing the UGC-CSIR-JRF (NTA Ref. No: 191620004555) fellowship to KC. This work is supported by the funding agency **ANRF (erstwhile SERB), India** (Grant no. **CRG/2023/002165**).

