## Supplementary material - Text, Figures and tables for "Deciphering the dynamical origin of the Adipogenic commitment during stress-induced disruption of the circadian rhythm"

**Construction of mathematical model**

To investigate the influence of circadian rhythms on adipogenic differentiation, we developed an integrated mathematical model comprising a circadian clock module and an adipogenic differentiation module (**Fig. 1D**). The circadian module captures the essential negative feedback loops responsible for the generation of self-sustained oscillations, whereas the adipogenic module describes the molecular interactions governing adipocyte differentiation. The two modules are coupled through glucocorticoid (GC) signalling, allowing the circadian clock to dynamically regulate adipogenic progression.

The model consists of 14 ordinary differential equations (ODEs) describing the temporal evolution of mRNAs and proteins involved in the circadian and adipogenic regulatory networks (**Table S2**). Transcription, translation, activation, inactivation and degradation processes were modelled using ODEs based on mass-action kinetics, Michaelis-Menten kinetics, and Hill functions wherever appropriate. Scaling factors were incorporated to convert concentration-based quantities into molecular abundances and to represent fold-change perturbations in different molecular species.

**Circadian Network**

The circadian network consists of three interconnected negative feedback loops. The first feedback loop (Module 1) corresponds to the core molecular clock formed by Bmal1 and PerCry complex. The second feedback loop is mediated through glucocorticoid signalling, while the third feedback loop involves Rev mediated repression of Bmal1.

The dynamics of Bmal1 (Bmal1-Clock complex) are described by **Eq. 1**. Bmal1 synthesis is negatively regulated by both PerCry complex and Rev protein through inhibitory saturation functions (1^st^ and 2^nd^ terms in **Eq. 1** respectively). These inhibitory interactions capture the transcriptional repression exerted by PerCry complex and Rev protein on Bmal1 expression. Bmal1 undergoes first-order degradation represented by the final term in **Eq. 1**.

$\frac{dBmal1}{dt}=sfcirc⸱x⸱\left( \frac{k_{sbmal1p}}{J_{bmal1p}+\frac{PerCry}{sfcirc⸱k_{mpercry}}}+\frac{k_{sbmal1r}}{J_{bmal1r}+\frac{Rev}{sfcirc⸱sfrev⸱k_{mrev}}} \right)-x⸱k_{dbmal1}⸱Bmal1$	**Eq. 1**

The dynamics of Per mRNA are described by **Eq. 2**. Per transcription consists of three components: a basal synthesis rate (1^st^ term in **Eq. 2**), activation by Bmal1 expression (2^nd^ term in **Eq. 2**) and activation by glucocorticoid signalling (3^rd^ term in **Eq. 2**). The degradation of Per mRNA is modelled as a first-order process (last term in **Eq. 2**).

$$\frac{dmPer}{dt}=sfcirc⸱x⸱b_{mper}+x⸱k_{smper}⸱Bmal1+x⸱k_{GC}⸱GC-x⸱k_{dmper}⸱mPer$$

**Eq. 2**

The dynamics of Per protein and the PerCry complex are represented by **Eq. 3** and **Eq. 4** respectively. Per protein is synthesized from Per mRNA (1^st^ term in **Eq. 3**) and participates in reversible complex formation with constitutively available Cry protein (2^nd^ & 3^rd^ terms in **Eq. 3** and 1^st^ & 2^nd^ terms in **Eq. 4**). Explicit Cry dynamics are not considered in the present model; instead, the total PerCry complex is represented as a single variable. Per protein and PerCry complex undergo both linear degradation (4^th^ term in **Eq. 3** and 3^rd^ term in **Eq. 4** respectively) and saturable degradation processes (last terms in **Eq. 3** and **Eq. 4** respectively). Since Bmal1 activates Per expression through Per mRNA (**Eq. 2** and **Eq. 3**), while PerCry complex represses Bmal1 synthesis (**Eq. 1**), this interaction forms the central circadian negative feedback loop. The reversible conversion between Per and PerCry introduces an effective delay in the negative feedback loop, which is essential for the generation of sustained circadian oscillations.

$\frac{dPer}{dt}=sfper⸱x⸱k_{sper}⸱mPer-x⸱k_{a}⸱Per+sfper⸱x⸱k_{d}⸱PerCry-x⸱k_{dper}⸱Per-\frac{x⸱k_{per}⸱Per}{J+\frac{q⸱Per}{sfcirc⸱sfper}+\frac{r⸱PerCry}{sfcirc}}$	**Eq. 3**

$\frac{dPerCry}{dt}=\frac{x⸱k_{a}⸱Per}{sfper}-x⸱k_{d}⸱PerCry-x⸱k_{dpercry}⸱PerCry-\frac{x⸱k_{percry}⸱PerCry}{J+\frac{q⸱Per}{sfcirc⸱sfper}+\frac{r⸱PerCry}{sfcirc}}$

**Eq. 4**

The second circadian feedback loop (Module 2) involves glucocorticoid signalling. The dynamics of GC are described in **Eq. 5**. GC is synthesized through a basal production term (1^st^ term in **Eq. 5**) and an additional regulated synthesis term which is inhibited by Per protein through a Hill-type repression function (2^nd^ term in **Eq. 5**). GC degradation follows first-order kinetics (last term in **Eq. 5**). Since GC promotes Per transcription (**Eq. 2**), whereas Per suppresses GC production (**Eq. 5**), these interactions establish the second negative feedback loop.

$\frac{dGC}{dt}=sfcirc⸱x⸱b_{GC}+\frac{sfcirc⸱x⸱k_{sGC}}{J_{GC}^{m}+\left( \frac{Per}{sfcirc⸱sfper⸱k_{mGC}} \right)^{m}}-x⸱k_{dGC}⸱GC$	**Eq. 5**

The third negative feedback loop (Module 3) is mediated by Rev. The dynamics of Rev mRNA and protein are represented by **Eq. 6** and **Eq. 7** respectively. Rev transcription is driven by basal synthesis (1^st^ term in **Eq. 6**), Bmal1 mediated synthesis (2^nd^ term in **Eq. 6**) and GC dependent activation (3^rd^ term in **Eq. 6**), while degradation occurs through a first-order process (last term in **Eq. 6**). Rev protein synthesis modelled using a Hill function to account for nonlinear regulation associated with mRNA translation (1^st^ term in **Eq. 7**). Rev degradation is represented using a saturable degradation term (last term in **Eq. 7**). Since Rev suppresses Bmal1 synthesis (**Eq. 1**), while Bmal1 activates Rev expression (**Eq. 6**), these interactions form the third negative feedback loop within the circadian regulatory network.

$\frac{dmRev}{dt}=sfcirc⸱sfrev1⸱x⸱b_{mrev}+sfrev1⸱x⸱k_{smrev}⸱Bmal1+sfrev1⸱x⸱k_{rGC}⸱GC-x⸱k_{dmrev}⸱mRev$	**Eq. 6**

$\frac{dRev}{dt}=sfsirc⸱sfrev⸱x⸱k_{srev}⸱\left( \frac{mRev^{h}}{\left( sfcirc⸱sfrev1⸱J_{rev} \right)^{h}+mRev^{h}} \right)-sfcirc⸱sfrev⸱x⸱k_{drev}⸱\left( \frac{Rev}{\left( sfcirc⸱sfrev⸱J_{drev} \right)+Rev} \right)$	**Eq. 7**

**Adipogenic Network**

The adipogenic module consists of three major molecular regulators: PPARγ, CEBPA and FABP4. The model incorporates two interconnected positive feedback loops: one between PPARγ and CEBPA, and another between PPARγ and FABP4.

The dynamics of PPARγ mRNA are described by Eq. 8. PPARγ transcription consists of a basal synthesis term (1^st^ term in **Eq. 8**) together with activation terms mediated by CEBPA (2^nd^ term in **Eq. 8**) and FABP4 (3^rd^ term in **Eq. 8**). Both activation processes are represented using Hill functions to account for cooperative transcriptional regulation. PPARγ mRNA degradation follows first-order kinetics (last term in **Eq. 8**).

$\frac{dmPPAR\gamma}{dt}=sf⸱sf1⸱sfP⸱sfP1⸱y⸱\left( k_{sP}+\frac{k_{P1}⸱CEBPA^{n}}{\left( sf⸱sfC⸱K_{mP1} \right)^{n}+CEBPA^{n}}+\frac{k_{P2}⸱FABP4^{n}}{\left( sf⸱sfF⸱K_{mP2} \right)^{n}+FABP4^{n}} \right)-y⸱k_{dmP}⸱mPPAR\gamma$	**Eq. 8**

To account for post-translational regulation, PPARγ protein is assumed to exist in inactive and active forms. The dynamics of inactive and active PPARγ are represented by **Eq. 9** and **Eq. 10** respectively. Inactive PPARγ is synthesized through translation of PPARγ mRNA (1^st^ term in **Eq. 9**), converted into the active form through a stimulus-independent and a stimulus-dependent activation processes (3^rd^ and 4^th^ term in **Eq. 9** respectively), regenerated through inactivation of active PPARγ (2^nd^ term in **Eq. 9**), and degraded through first-order kinetics (last term in **Eq. 9**). Active PPARγ is generated from the inactive form (1^st^ and 2^nd^ term in **Eq. 10**), can revert back to the inactive form (3^rd^ term in **Eq. 10**) and undergoes first-order degradation (last term in **Eq. 10**). The activation rate is regulated by an external stimulus duration term that control adipogenic induction.

$\frac{dPPAR\gamma_{i}}{dt}=\frac{y⸱k_{tP}⸱mPPAR\gamma}{sf1⸱sfP1}+y⸱k_{inact}⸱PPAR\gamma_{a}-y⸱\left( k_{0}+k_{act}^{'}⸱Stimulus \right)⸱PPAR\gamma_{i}-y⸱k_{dP}⸱PPAR\gamma_{i}$	**Eq. 9**

$\frac{dPPAR\gamma_{a}}{dt}=y⸱\left( k_{0}+k_{act}^{'}⸱Stimulus \right)⸱PPAR\gamma_{i}-y⸱k_{inact}⸱PPAR\gamma_{a}-y⸱k_{dP}⸱PPAR\gamma_{a}$	**Eq. 10**

The dynamics of CEBPA mRNA and protein are described by **Eq. 11** and **Eq. 12** respectively. CEBPA mRNA synthesis consists of a basal production term (1^st^ term in **Eq. 11**) and a PPARγ mediated activation term represented by a Hill function (2^nd^ term in **Eq. 11**). Its degradation follows first-order kinetics (last term in **Eq. 11**). The corresponding protein is synthesized through translation (1^st^ term in **Eq. 12**) and degraded through first-order kinetics (last term in **Eq. 12**). Since active PPARγ activates CEBPA expression (**Eq. 11**) and CEBPA in turn enhances PPARγ transcription (**Eq. 8**), these interactions constitute the first positive feedback loop of the adipogenic network.

$\frac{dmCEBPA}{dt}=sf⸱sf1⸱sfC⸱sfC1⸱y⸱\left( b_{mCEBPA}+\frac{k_{C}⸱PPAR\gamma_{a}^{n}}{\left( sf⸱sfP⸱K_{mC} \right)^{n}+PPAR\gamma_{a}^{n}} \right)-y⸱k_{dmC}⸱mCEBPA$	**Eq. 11**

$\frac{dCEBPA}{dt}=\frac{y⸱k_{tC}⸱mCEBPA}{sf1⸱sfC1}-y⸱k_{dC}⸱CEBPA$	**Eq. 12**

The dynamics of FABP4 mRNA and protein are described by **Eq. 13** and **Eq. 14** respectively. FABP4 mRNA synthesis consists of a basal production term (1^st^ term in **Eq. 13**) and a PPARγ mediated activation term represented by a Hill function (2^nd^ term in **Eq. 13**). The corresponding protein is synthesized through translation (1^st^ term in **Eq. 14**), while both mRNA and protein species undergo first-order degradation (last terms in **Eq. 13** and **Eq. 14** respectively). FABP4 positively regulates PPARγ transcription (**Eq. 8**), thereby forming the second positive feedback loop within the adipogenic module.

$\frac{dmFABP4}{dt}=sf⸱sf1⸱sfF⸱sfF1⸱y⸱\left( b_{mFABP4}+\frac{k_{F}⸱PPAR\gamma_{a}^{n}}{\left( sf⸱sfP⸱K_{mF} \right)^{n}+PPAR\gamma_{a}^{n}} \right)-y⸱k_{dmF}⸱mFABP4$	**Eq. 13**

$\frac{dFABP4}{dt}=\frac{y⸱k_{tF}⸱mFABP4}{sf1⸱sfF1}-y⸱k_{dF}⸱FABP4$	**Eq. 14**

The combined action of these two positive feedback loops enables switch-like activation of adipogenic differentiation and stabilizes the differentiated adipocyte phenotype once the activation threshold has been crossed.

In the integrated model, the circadian clock generates oscillatory GC dynamics through the Per-GC negative feedback loop. The GC signal acts as an upstream regulator of adipogenic induction (**Table S5**).

In simulations involving only the adipogenic module (**Fig. 1B** and **Table S1**), the stimulus term was represented either as a constant signal or as a pulsatile input modelled using a Heaviside function (**Materials and methods**). In contrast, within the integrated model, the stimulus is dynamically controlled by the circadian GC signal, allowing the investigation of how circadian oscillatory hormone profiles, and clock perturbations influence adipogenic commitment.

| Table S1. Ordinary Differential Equations for the Adipogenic network in Fig. 1B |  |
| --- | --- |
| $\frac{\boldsymbol{dmPPAR\gamma}}{\mathbf{dt}}\mathbf{=sf}\mathbf{⸳}\mathbf{sf1}\mathbf{⸳}\mathbf{sfP}\mathbf{⸳}\mathbf{sfP1}\mathbf{⸳}\mathbf{y}\mathbf{⸳}\left( \mathbf{k}_{\mathbf{sP}}\mathbf{+}\frac{\mathbf{k}_{\mathbf{P1}}\mathbf{⸳}\mathbf{CEBP}\mathbf{A}^{\mathbf{n}}}{\left( \mathbf{sf}\mathbf{⸳}\mathbf{sfC}\mathbf{⸳}\mathbf{K}_{\mathbf{mP1}} \right)^{\mathbf{n}}\mathbf{+CEBP}\mathbf{A}^{\mathbf{n}}}\mathbf{+}\frac{\mathbf{k}_{\mathbf{P2}}\mathbf{⸳}\mathbf{FABP}\mathbf{4}^{\mathbf{n}}}{\left( \mathbf{sf}\mathbf{⸳}\mathbf{sfF}\mathbf{⸳}\mathbf{K}_{\mathbf{mP2}} \right)^{\mathbf{n}}\mathbf{+FABP}\mathbf{4}^{\mathbf{n}}} \right)\mathbf{-y}\mathbf{⸳}\mathbf{k}_{\mathbf{dmP}}\mathbf{⸳}\boldsymbol{mPPAR\gamma}$ | **1** |
| $\frac{\mathbf{dPPAR}\boldsymbol{\gamma}_{\mathbf{i}}}{\mathbf{dt}}\mathbf{=}\frac{\mathbf{y}\mathbf{⸳}\mathbf{k}_{\mathbf{tP}}\mathbf{⸳}\boldsymbol{mPPAR\gamma}}{\mathbf{sf1}\mathbf{⸳}\mathbf{sfP1}}\mathbf{+y}\mathbf{⸳}\mathbf{k}_{\mathbf{inact}}\mathbf{⸳}\mathbf{PPAR}\boldsymbol{\gamma}_{\mathbf{a}}\mathbf{-y}\mathbf{⸳}\left( \mathbf{k}_{\mathbf{0}}\mathbf{+}\mathbf{k'}_{\mathbf{act}}\mathbf{⸳}\mathbf{Stimulus} \right)\mathbf{⸳}\mathbf{PPAR}\boldsymbol{\gamma}_{\mathbf{i}}\mathbf{-y}\mathbf{⸳}\mathbf{k}_{\mathbf{dP}}\mathbf{⸳}\mathbf{PPAR}\boldsymbol{\gamma}_{\mathbf{i}}$ | **2** |
| $\frac{\mathbf{dPPAR}\boldsymbol{\gamma}_{\mathbf{a}}}{\mathbf{dt}}\mathbf{=y}\mathbf{⸳}\left( \mathbf{k}_{\mathbf{0}}\mathbf{+}\mathbf{k'}_{\mathbf{act}}\mathbf{⸳}\mathbf{Stimulus} \right)\mathbf{⸳}\mathbf{PPAR}\boldsymbol{\gamma}_{\mathbf{i}}\mathbf{-y}\mathbf{⸳}\mathbf{k}_{\mathbf{inact}}\mathbf{⸳}\mathbf{PPAR}\boldsymbol{\gamma}_{\mathbf{a}}\mathbf{-y}\mathbf{⸳}\mathbf{k}_{\mathbf{dP}}\mathbf{⸳}\mathbf{PPAR}\boldsymbol{\gamma}_{\mathbf{a}}$ | **3** |
| $\frac{\mathbf{dmCEBPA}}{\mathbf{dt}}\mathbf{=sf}\mathbf{⸳}\mathbf{sf1}\mathbf{⸳}\mathbf{sfC}\mathbf{⸳}\mathbf{sfC1}\mathbf{⸳}\mathbf{y}\mathbf{⸳}\left( \mathbf{b}_{\mathbf{mCEBPA}}\mathbf{+}\frac{\mathbf{k}_{\mathbf{C}}\mathbf{⸳}\mathbf{PPAR}\boldsymbol{\gamma}_{\mathbf{a}}^{\mathbf{n}}}{\left( \mathbf{sf}\mathbf{⸳}\mathbf{sfP}\mathbf{⸳}\mathbf{K}_{\mathbf{mC}} \right)^{\mathbf{n}}\mathbf{+PPAR}\boldsymbol{\gamma}_{\mathbf{a}}^{\mathbf{n}}} \right)\mathbf{-y}\mathbf{⸳}\mathbf{k}_{\mathbf{dmC}}\mathbf{⸳}\mathbf{mCEBPA}$ | **4** |
| $\frac{\mathbf{dCEBPA}}{\mathbf{dt}}\mathbf{=}\frac{\mathbf{y}\mathbf{⸳}\mathbf{k}_{\mathbf{tC}}\mathbf{⸳}\mathbf{mCEBPA}}{\mathbf{sf1}\mathbf{⸳}\mathbf{sfC1}}\mathbf{-y}\mathbf{⸳}\mathbf{k}_{\mathbf{dC}}\mathbf{⸳}\mathbf{CEBPA}$ | **5** |
| $\frac{\mathbf{dmFABP4}}{\mathbf{dt}}\mathbf{=sf}\mathbf{⸳}\mathbf{sf1}\mathbf{⸳}\mathbf{sfF}\mathbf{⸳}\mathbf{sfF1}\mathbf{⸳}\mathbf{y}\mathbf{⸳}\left( \mathbf{b}_{\mathbf{mFABP4}}\mathbf{+}\frac{\mathbf{k}_{\mathbf{F}}\mathbf{⸳}\mathbf{PPAR}\boldsymbol{\gamma}_{\mathbf{a}}^{\mathbf{n}}}{\left( \mathbf{sf}\mathbf{⸳}\mathbf{sfF}\mathbf{⸳}\mathbf{K}_{\mathbf{mF}} \right)^{\mathbf{n}}\mathbf{+PPAR}\boldsymbol{\gamma}_{\mathbf{a}}^{\mathbf{n}}} \right)\mathbf{-y}\mathbf{⸳}\mathbf{k}_{\mathbf{dmF}}\mathbf{⸳}\mathbf{mFABP4}$ | **6** |
| $\frac{\mathbf{dFABP4}}{\mathbf{dt}}\mathbf{=}\frac{\mathbf{y}\mathbf{⸳}\mathbf{k}_{\mathbf{tF}}\mathbf{⸳}\mathbf{mFABP4}}{\mathbf{sf1}\mathbf{⸳}\mathbf{sfF1}}\mathbf{-y}\mathbf{⸳}\mathbf{k}_{\mathbf{dF}}\mathbf{⸳}\mathbf{FABP4}$ | **7** |

| Table S2. Ordinary Differential Equations for the integrated model network in Fig. 1D |  |
| --- | --- |
| $\frac{\mathbf{dBmal1}}{\mathbf{dt}}\mathbf{=sfcirc}\mathbf{⸳}\mathbf{x}\mathbf{⸳}\left( \frac{\mathbf{k}_{\mathbf{sbmal1p}}}{\mathbf{J}_{\mathbf{bmal1p}}\mathbf{+}\frac{\mathbf{PerCry}}{\mathbf{sfcirc}\mathbf{⸳}\mathbf{k}_{\mathbf{mpercry}}}}\mathbf{+}\frac{\mathbf{k}_{\mathbf{sbaml1r}}}{\mathbf{J}_{\mathbf{bmal1r}}\mathbf{+}\frac{\mathbf{Rev}}{\mathbf{sfcirc}\mathbf{⸳}\mathbf{sfrev}\mathbf{⸳}\mathbf{k}_{\mathbf{mrev}}}} \right)\mathbf{-x}\mathbf{⸳}\mathbf{k}_{\mathbf{dbmal1}}\mathbf{⸳}\mathbf{Bmal1}$ | **1** |
| $\frac{\mathbf{dmPer}}{\mathbf{dt}}\mathbf{=sfcirc}\mathbf{⸳}\mathbf{x}\mathbf{⸳}\mathbf{b}_{\mathbf{mper}}\mathbf{+x}\mathbf{⸳}\mathbf{k}_{\mathbf{smper}}\mathbf{⸳}\mathbf{Bmal1+x}\mathbf{⸳}\mathbf{k}_{\mathbf{GC}}\mathbf{⸳}\mathbf{GC-x}\mathbf{⸳}\mathbf{k}_{\mathbf{dmper}}\mathbf{⸳}\mathbf{mPer}$ | **2** |
| $\frac{\mathbf{dPer}}{\mathbf{dt}}\mathbf{=sfper}\mathbf{⸳}\mathbf{x}\mathbf{⸳}\mathbf{k}_{\mathbf{sper}}\mathbf{⸳}\mathbf{mPer-}{\mathbf{x}\mathbf{⸳}\mathbf{k}}_{\mathbf{a}}\mathbf{⸳}\mathbf{Per+sfper}\mathbf{⸳}\mathbf{x}\mathbf{⸳}\mathbf{k}_{\mathbf{d}}\mathbf{⸳}\mathbf{PerCry-x}\mathbf{⸳}\mathbf{k}_{\mathbf{dper}}\mathbf{⸳}\mathbf{Per-}\frac{\mathbf{x}\mathbf{⸳}\mathbf{k}_{\mathbf{per}}\mathbf{⸳}\mathbf{Per}}{\mathbf{J+}\frac{\mathbf{q}\mathbf{⸳}\mathbf{Per}}{\mathbf{sfcirc}\mathbf{⸳}\mathbf{sfper}}\mathbf{+}\frac{\mathbf{r}\mathbf{⸳}\mathbf{PerCry}}{\mathbf{sfcirc}}}$ | **3** |
| $\frac{\mathbf{dPerCry}}{\mathbf{dt}}\mathbf{=}\frac{\mathbf{x}\mathbf{⸳}\mathbf{k}_{\mathbf{a}}\mathbf{⸳}\mathbf{Per}}{\mathbf{sfper}}\boldsymbol{-}\mathbf{x}\mathbf{⸳}\mathbf{k}_{\mathbf{d}}\mathbf{⸳}\mathbf{PerCry-x}\mathbf{⸳}\mathbf{k}_{\mathbf{dpercry}}\mathbf{⸳}\mathbf{PerCry-}\frac{\mathbf{x}\mathbf{⸳}\mathbf{k}_{\mathbf{percry}}\mathbf{⸳}\mathbf{PerCry}}{\mathbf{J+}\frac{\mathbf{q}\mathbf{⸳}\mathbf{Per}}{\mathbf{sfcirc}\mathbf{⸳}\mathbf{sfper}}\mathbf{+}\frac{\mathbf{r}\mathbf{⸳}\mathbf{PerCry}}{\mathbf{sfcirc}}}$ | **4** |
| $\frac{\mathbf{dGC}}{\mathbf{dt}}\mathbf{=sfcirc}\mathbf{⸳}\mathbf{x}\mathbf{⸳}\mathbf{b}_{\mathbf{GC}}\mathbf{+}\frac{\mathbf{sfcirc}\mathbf{⸳}\mathbf{x}\mathbf{⸳}\mathbf{k}_{\mathbf{sGC}}}{\mathbf{J}_{\mathbf{GC}}^{\mathbf{m}}\mathbf{+}\left( \frac{\mathbf{Per}}{\mathbf{sfcirc}\mathbf{⸳}\mathbf{sfper}\mathbf{⸳}\mathbf{k}_{\mathbf{mGC}}} \right)^{\mathbf{m}}}\mathbf{-x}\mathbf{⸳}\mathbf{k}_{\mathbf{dGC}}\mathbf{⸳}\mathbf{GC}$ | **5** |
| $\frac{\mathbf{dmRev}}{\mathbf{dt}}\mathbf{=sfcirc}\mathbf{⸳}\mathbf{sfrev1}\mathbf{⸳}\mathbf{x}\mathbf{⸳}\mathbf{b}_{\mathbf{mrev}}\mathbf{+sfrev1}\mathbf{⸳}\mathbf{x}\mathbf{⸳}\mathbf{k}_{\mathbf{smrev}}\mathbf{⸳}\mathbf{Bmal1+sfrev1}\mathbf{⸳}\mathbf{x}\mathbf{⸳}\mathbf{k}_{\mathbf{rGC}}\mathbf{⸳}\mathbf{GC-x}\mathbf{⸳}\mathbf{k}_{\mathbf{dmrev}}\mathbf{⸳}\mathbf{mRev}$ | **6** |
| $\frac{\mathbf{dRev}}{\mathbf{dt}}\mathbf{=sfcirc}\mathbf{⸳}\mathbf{sfrev}\mathbf{⸳}\mathbf{x}\mathbf{⸳}\mathbf{k}_{\mathbf{srev}}\mathbf{⸳}\left( \frac{\mathbf{mRe}\mathbf{v}^{\mathbf{h}}}{\left( \mathbf{sfcirc}\mathbf{⸳}\mathbf{sfrev1}\mathbf{⸳}\mathbf{J}_{\mathbf{rev}} \right)^{\mathbf{h}}\mathbf{+mRe}\mathbf{v}^{\mathbf{h}}} \right)\boldsymbol{-}\mathbf{sfcirc}\mathbf{⸳}\mathbf{sfrev}\mathbf{⸳}\mathbf{x}\mathbf{⸳}\mathbf{k}_{\mathbf{drev}}\mathbf{⸳}\left( \frac{\mathbf{Rev}}{\left( \mathbf{sfcirc}\mathbf{⸳}\mathbf{sfrev}\mathbf{⸳}\mathbf{J}_{\mathbf{drev}} \right)\mathbf{+Rev}} \right)$ | **7** |
| $\frac{\boldsymbol{dmPPAR\gamma}}{\mathbf{dt}}\mathbf{=sf}\mathbf{⸳}\mathbf{sf1}\mathbf{⸳}\mathbf{sfP}\mathbf{⸳}\mathbf{sfP1}\mathbf{⸳}\mathbf{y}\mathbf{⸳}\left( \mathbf{k}_{\mathbf{sP}}\mathbf{+}\frac{\mathbf{k}_{\mathbf{P1}}\mathbf{⸳}\mathbf{CEBP}\mathbf{A}^{\mathbf{n}}}{\left( \mathbf{sf}\mathbf{⸳}\mathbf{sfC}\mathbf{⸳}\mathbf{K}_{\mathbf{mP1}} \right)^{\mathbf{n}}\mathbf{+CEBP}\mathbf{A}^{\mathbf{n}}}\mathbf{+}\frac{\mathbf{k}_{\mathbf{P2}}\mathbf{⸳}\mathbf{FABP}\mathbf{4}^{\mathbf{n}}}{\left( \mathbf{sf}\mathbf{⸳}\mathbf{sfF}\mathbf{⸳}\mathbf{K}_{\mathbf{mP2}} \right)^{\mathbf{n}}\mathbf{+FABP}\mathbf{4}^{\mathbf{n}}} \right)\mathbf{-y}\mathbf{⸳}\mathbf{k}_{\mathbf{dmP}}\mathbf{⸳}\boldsymbol{mPPAR\gamma}$ | **8** |
| $\frac{\mathbf{dPPAR}\boldsymbol{\gamma}_{\mathbf{i}}}{\mathbf{dt}}\mathbf{=}\frac{\mathbf{y}\mathbf{⸳}\mathbf{k}_{\mathbf{tP}}\mathbf{⸳}\boldsymbol{mPPAR\gamma}}{\mathbf{sf1}\mathbf{⸳}\mathbf{sfP1}}\mathbf{+y}\mathbf{⸳}\mathbf{k}_{\mathbf{inact}}\mathbf{⸳}\mathbf{PPAR}\boldsymbol{\gamma}_{\mathbf{a}}\mathbf{-y}\mathbf{⸳}\left( \mathbf{k}_{\mathbf{0}}\mathbf{+}\mathbf{k'}_{\mathbf{act}}\mathbf{⸳}\mathbf{Stimulus} \right)\mathbf{⸳}\mathbf{PPAR}\boldsymbol{\gamma}_{\mathbf{i}}\mathbf{-y}\mathbf{⸳}\mathbf{k}_{\mathbf{dP}}\mathbf{⸳}\mathbf{PPAR}\boldsymbol{\gamma}_{\mathbf{i}}$ | **9** |
| $\frac{\mathbf{dPPAR}\boldsymbol{\gamma}_{\mathbf{a}}}{\mathbf{dt}}\mathbf{=y}\mathbf{⸳}\left( \mathbf{k}_{\mathbf{0}}\mathbf{+}\mathbf{k'}_{\mathbf{act}}\mathbf{⸳}\mathbf{Stimulus} \right)\mathbf{⸳}\mathbf{PPAR}\boldsymbol{\gamma}_{\mathbf{i}}\mathbf{-y}\mathbf{⸳}\mathbf{k}_{\mathbf{inact}}\mathbf{⸳}\mathbf{PPAR}\boldsymbol{\gamma}_{\mathbf{a}}\mathbf{-y}\mathbf{⸳}\mathbf{k}_{\mathbf{dP}}\mathbf{⸳}\mathbf{PPAR}\boldsymbol{\gamma}_{\mathbf{a}}$ | **10** |
| $\frac{\mathbf{dmCEBPA}}{\mathbf{dt}}\mathbf{=sf}\mathbf{⸳}\mathbf{sf1}\mathbf{⸳}\mathbf{sfC}\mathbf{⸳}\mathbf{sfC1}\mathbf{⸳}\mathbf{y}\mathbf{⸳}\left( \mathbf{b}_{\mathbf{mCEBPA}}\mathbf{+}\frac{\mathbf{k}_{\mathbf{C}}\mathbf{⸳}\mathbf{PPAR}\boldsymbol{\gamma}_{\mathbf{a}}^{\mathbf{n}}}{\left( \mathbf{sf}\mathbf{⸳}\mathbf{sfP}\mathbf{⸳}\mathbf{K}_{\mathbf{mC}} \right)^{\mathbf{n}}\mathbf{+PPAR}\boldsymbol{\gamma}_{\mathbf{a}}^{\mathbf{n}}} \right)\mathbf{-y}\mathbf{⸳}\mathbf{k}_{\mathbf{dmC}}\mathbf{⸳}\mathbf{mCEBPA}$ | **11** |
| $\frac{\mathbf{dCEBPA}}{\mathbf{dt}}\mathbf{=}\frac{\mathbf{y}\mathbf{⸳}\mathbf{k}_{\mathbf{tC}}\mathbf{⸳}\mathbf{mCEBPA}}{\mathbf{sf1}\mathbf{⸳}\mathbf{sfC1}}\mathbf{-y}\mathbf{⸳}\mathbf{k}_{\mathbf{dC}}\mathbf{⸳}\mathbf{CEBPA}$ | **12** |
| $\frac{\mathbf{dmFABP4}}{\mathbf{dt}}\mathbf{=sf}\mathbf{⸳}\mathbf{sf1}\mathbf{⸳}\mathbf{sfF}\mathbf{⸳}\mathbf{sfF1}\mathbf{⸳}\mathbf{y}\mathbf{⸳}\left( \mathbf{b}_{\mathbf{mFABP4}}\mathbf{+}\frac{\mathbf{k}_{\mathbf{F}}\mathbf{⸳}\mathbf{PPAR}\boldsymbol{\gamma}_{\mathbf{a}}^{\mathbf{n}}}{\left( \mathbf{sf}\mathbf{⸳}\mathbf{sfF}\mathbf{⸳}\mathbf{K}_{\mathbf{mF}} \right)^{\mathbf{n}}\mathbf{+PPAR}\boldsymbol{\gamma}_{\mathbf{a}}^{\mathbf{n}}} \right)\mathbf{-y}\mathbf{⸳}\mathbf{k}_{\mathbf{dmF}}\mathbf{⸳}\mathbf{mFABP4}$ | **13** |
| $\frac{\mathbf{dFABP4}}{\mathbf{dt}}\mathbf{=}\frac{\mathbf{y}\mathbf{⸳}\mathbf{k}_{\mathbf{tF}}\mathbf{⸳}\mathbf{mFABP4}}{\mathbf{sf1}\mathbf{⸳}\mathbf{sfF1}}\mathbf{-y}\mathbf{⸳}\mathbf{k}_{\mathbf{dF}}\mathbf{⸳}\mathbf{FABP4}$ | **14** |

**Table S3.** Abbreviated names of the variables and their description

| Abbreviated names | Description |
| --- | --- |
| Bmal1 | **Bmal1 ̶ Clock complex** |
| mPer | **Per mRNA** |
| Per | **Per protein** |
| PerCry | **Per ̶ Cry complex** |
| GC | **Glucocorticoids** |
| mRev | **Rev mRNA** |
| Rev | **Rev protein** |
| mPPAR$\boldsymbol{\gamma}$ | **PPAR**$\boldsymbol{\gamma}$ **mRNA** |
| PPAR$\boldsymbol{\gamma}$_i_ | **Inactivated PPAR**$\boldsymbol{\gamma}$ **protein** |
| PPAR$\boldsymbol{\gamma}$_a_ | **Activated PPAR**$\boldsymbol{\gamma}$ **protein** |
| mCEBPA | **CEBPA mRNA** |
| CEBPA | **CEBPA protein** |
| mFABP4 | **FABP4 mRNA** |
| FABP4 | **FABP4 protein** |

| **Table S4.** Reaction governing the integrated model network in Fig. 1D | | | |
| --- | --- | --- | --- |
| **Sl. No.** | **Reaction** | **Kinetics** | **Description** |
| **Bmal1 ̶ Clock complex** | | | |
| R1 | $PerCry ⟞ Bmal1$ | $\frac{\mathrm{sfcirc}⸳x⸳k_{sbmal1p}}{J_{bmal1p}+\frac{\mathrm{PerCry}}{\mathrm{sfcirc}⸳k_{\mathrm{mpercry}}}}$ | Synthesis of Bmal1 ̶ Clock complex regulated through Per ̶ Cry pathway |
| R2 | $Rev ⟞ Bmal1$ | $\frac{\mathrm{sfcirc}⸳x⸳k_{sbmal1r}}{J_{bmal1r}+\frac{\mathrm{Rev}}{\mathrm{sfcirc}⸳\mathrm{sfrev}⸳k_{\mathrm{mrev}}}}$ | Synthesis of Bmal1 ̶ Clock complex regulated through Rev pathway |
| R3 | $Bmal1\to\therefore$ | $x⸳k_{dbmal1}⸳Bmal1$ | Degradation of Bmal1 ̶ Clock complex |
| **Per mRNA** | | | |
| R4 | $\to mPer$ | $\mathrm{sfcirc}⸳x⸳b_{\mathrm{mper}}$ | Basal transcription of Per mRNA |
| R5 | $\underset{\to}{Bmal1}mPer$ | $x⸳k_{\mathrm{smper}}⸳Bmal1$ | Bmal1 ̶ Clock complex mediated transcription of Per mRNA |
| R6 | $\underset{\to}{GC}mPer$ | $x⸳k_{\mathrm{GC}}⸳\mathrm{GC}$ | GC mediated transcription of Per mRNA |
| R7 | $mPer\to\therefore$ | $x⸳k_{\mathrm{dmper}}⸳\mathrm{mPer}$ | Degradation of Per mRNA |
| **Per protein** | | | |
| R8 | $\underset{\to}{mPer}Per$ | $\mathrm{sfper}⸳x⸳k_{\mathrm{sper}}⸳\mathrm{mPer}$ | Translation of Per protein |
| R9 | $PerCry\to Per$ | $x⸳k_{d}⸳\mathrm{PerCry}$ | Dissociation of Per ̶ Cry complex |
| R10 | $Per\to\therefore$ | $x⸳k_{\mathrm{dper}}⸳\mathrm{Per}$ | Degradation of Per protein |
| R11 | $Per\underset{\to}{PerCry} \therefore$ | $\frac{x⸳k_{\mathrm{per}}⸳\mathrm{Per}}{J+\frac{q⸳\mathrm{Per}}{\mathrm{sfcirc}⸳\mathrm{sfper}}+\frac{r⸳\mathrm{PerCry}}{\mathrm{sfcirc}}}$ | Non-linear degradation of Per protein |
| **Per ̶ Cry complex** | | | |
| R12 | $Per\to PerCry$ | $\frac{x⸳k_{a}⸳\mathrm{Per}}{\mathrm{sfper}}$ | Association of Per ̶ Cry complex |
| R13 | $PerCry\to\therefore$ | $x⸳k_{\mathrm{dpercry}}⸳\mathrm{PerCry}$ | Degradation of Per ̶ Cry complex |
| R14 | $PerCry\underset{\to}{Per} \therefore$ | $\frac{x⸳k_{\mathrm{percry}}⸳\mathrm{PerCry}}{J+\frac{q⸳\mathrm{Per}}{\mathrm{sfcirc}⸳\mathrm{sfper}}+\frac{r⸳\mathrm{PerCry}}{\mathrm{sfcirc}}}$ | Non-linear degradation of Per ̶ Cry complex |
| **GC** | | | |
| R15 | $\to GC$ | $\mathrm{sfcirc}⸳x⸳b_{\mathrm{GC}}$ | Basal synthesis of GC |
| R16 | $Per ⟞ GC$ | $\frac{\mathrm{sfcirc}⸳x⸳k_{\mathrm{sGC}}}{J_{\mathrm{GC}}^{m}+\left( \frac{\mathrm{Per}}{\mathrm{sfcirc}⸳\mathrm{sfper}⸳k_{\mathrm{mGC}}} \right)^{m}}$ | Synthesis of GC through Per pathway |
| R17 | $GC\to\therefore$ | $x⸳k_{\mathrm{dGC}}⸳\mathrm{GC}$ | Degradation of GC |
| **Rev mRNA** | | | |
| R18 | $\to mRev$ | $\mathrm{sfcirc}⸳sfrev1⸳x⸳b_{\mathrm{mrev}}$ | Basal transcription of Rev mRNA |
| R19 | $\underset{\to}{Bmal1}mRev$ | $sfrev1⸳x⸳k_{\mathrm{smrev}}⸳Bmal1$ | Bmal1 ̶ Clock complex mediated transcription of Rev mRNA |
| R20 | $\underset{\to}{GC}mRev$ | $sfrev1⸳x⸳k_{\mathrm{rGC}}⸳\mathrm{GC}$ | GC mediated transcription of Rev mRNA |
| R21 | $mRev\to\therefore$ | $x⸳k_{\mathrm{dmrev}}⸳\mathrm{mRev}$ | Degradation of Rev mRNA |
| **Rev protein** | | | |
| R22 | $\underset{\to}{mRev}Rev$ | $\frac{\mathrm{sfcirc}⸳\mathrm{sfrev}⸳x⸳k_{\mathrm{srev}}⸳\mathrm{mRe}v^{h}}{\left( \mathrm{sfcirc}⸳sfrev1⸳J_{\mathrm{rev}} \right)^{h}+mRev^{h}}$ | Translation of Rev protein |
| R23 | $Rev\to\therefore$ | $\frac{\mathrm{sfcirc}⸳\mathrm{sfrev}⸳x⸳k_{\mathrm{drev}}⸳\mathrm{Rev}}{\left( \mathrm{sfcirc}⸳\mathrm{sfrev}⸳J_{\mathrm{drev}} \right)+Rev}$ | Degradation of Rev protein |
| **PPAR**$\boldsymbol{\gamma}$ **mRNA** | | | |
| R24 | $\to mPPAR\gamma$ | $\mathrm{sf}⸳sf1⸳\mathrm{sfP}⸳sfP1⸳y⸳k_{\mathrm{sp}}$ | Basal transcription of PPAR$\gamma$ mRNA |
| R25 | $\underset{\to}{CEBPA}mPPAR\gamma$ | $\frac{\mathrm{sf}⸳sf1⸳\mathrm{sfP}⸳sfP1⸳y⸳k_{P1}⸳\mathrm{CEBP}A^{n}}{\left( \mathrm{sf}⸳\mathrm{sfC}⸳K_{mP1} \right)^{n}+CEBPA^{n}}$ | CEBPA mediated transcription of PPAR$\gamma$ mRNA |
| R26 | $\underset{\to}{FABP4}mPPAR\gamma$ | $\frac{\mathrm{sf}⸳sf1⸳\mathrm{sfP}⸳sfP1⸳y⸳k_{P2}⸳\mathrm{FABP}4^{n}}{\left( \mathrm{sf}⸳\mathrm{sfF}⸳K_{mP2} \right)^{n}+FABP4^{n}}$ | FABP4 mediated transcription of PPAR$\gamma$ mRNA |
| R27 | $mPPAR\gamma\to\therefore$ | $y⸳k_{\mathrm{dmP}}⸳mPPAR\gamma$ | Degradation of PPAR$\gamma$ mRNA |
| **Inactivated PPAR**$\boldsymbol{\gamma}$ **protein** | | | |
| R28 | $\underset{\to}{mPPAR\gamma}PPAR\gamma_{i}$ | $\frac{y⸳k_{\mathrm{tP}}⸳mPPAR\gamma}{sf1⸳sfP1}$ | Translation of PPAR$\gamma$ protein |
| R29 | $PPAR\gamma_{a}\to PPAR\gamma_{i}$ | $y⸳k_{\mathrm{inact}}⸳\mathrm{PPAR}\gamma_{a}$ | Inactivation of PPAR$\gamma$ protein |
| R30 | $PPAR\gamma_{i}\to\therefore$ | $y⸳k_{\mathrm{dP}}⸳\mathrm{PPAR}\gamma_{i}$ | Degradation of inactivated PPAR$\gamma$ protein |
| **Activated PPAR**$\boldsymbol{\gamma}$ **protein** | | | |
| R31 | $PPAR\gamma_{i}\to PPAR\gamma_{a}$ | $y⸳k_{0}⸳\mathrm{PPAR}\gamma_{i}$ | Stimulus-independent basal activation of PPAR$\gamma$ protein |
| R32 | $PPAR\gamma_{i}\underset{\to}{Stimulus}PPAR\gamma_{a}$ | $y⸳{k'}_{act}⸳\mathrm{Stimulus}⸳\mathrm{PPAR}\gamma_{i}$ | Stimulus-induced activation of PPAR$\gamma$ protein |
| R33 | $PPAR\gamma_{a}\to\therefore$ | $y⸳k_{\mathrm{dP}}⸳\mathrm{PPAR}\gamma_{a}$ | Degradation of activated PPAR$\gamma$ protein |
| **CEBPA mRNA** | | | |
| R34 | $\to mCEBPA$ | $\mathrm{sf}⸳sf1⸳\mathrm{sfC}⸳sfC1⸳y⸳b_{\mathrm{mCEBPA}}$ | Basal transcription of CEBPA mRNA |
| R35 | $\underset{\to}{PPAR\gamma_{a}}mCEBPA$ | $\frac{\mathrm{sf}⸳sf1⸳\mathrm{sfC}⸳sfC1⸳y⸳k_{C}⸳ {PPAR\gamma}_{a}^{n}}{\left( \mathrm{sf}⸳\mathrm{sfP}⸳K_{\mathrm{mC}} \right)^{n}+{PPAR\gamma}_{a}^{n}}$ | PPAR$\gamma$ mediated transcription of CEBPA mRNA |
| R36 | $mCEBPA\to\therefore$ | $y⸳k_{\mathrm{dmC}}⸳\mathrm{mCEBPA}$ | Degradation of CEBPA mRNA |
| **CEBPA protein** | | | |
| R37 | $\underset{\to}{mCEBPA}CEBPA$ | $\frac{y⸳k_{\mathrm{tC}}⸳\mathrm{mCEBPA}}{sf1⸳sfC1}$ | Translation of CEBPA protein |
| R38 | $CEBPA\to\therefore$ | $y⸳k_{\mathrm{dC}}⸳\mathrm{CEBPA}$ | Degradation of CEBPA protein |
| **FABP4 mRNA** | | | |
| R39 | $\to mFABP4$ | $\mathrm{sf}⸳sf1⸳\mathrm{sfF}⸳sfF1⸳y⸳b_{mFABP4}$ | Basal transcription of FABP4 mRNA |
| R40 | $\underset{\to}{PPAR\gamma_{a}}mFABP4$ | $\frac{\mathrm{sf}⸳sf1⸳\mathrm{sfF}⸳sfF1⸳y⸳k_{F}⸳ {PPAR\gamma}_{a}^{n}}{\left( \mathrm{sf}⸳\mathrm{sfP}⸳K_{\mathrm{mF}} \right)^{n}+{PPAR\gamma}_{a}^{n}}$ | PPAR$\gamma$ mediated transcription of FABP4 mRNA |
| R41 | $mFABP4\to\therefore$ | $y⸳k_{\mathrm{dmF}}⸳mFABP4$ | Degradation of FABP4 mRNA |
| **FABP4 protein** | | | |
| R42 | $\underset{\to}{mFABP4}FABP4$ | $\frac{y⸳k_{\mathrm{tF}}⸳mFABP4}{sf1⸳sfF1}$ | Translation of FABP4 protein |
| R43 | $FABP4\to\therefore$ | $y⸳k_{\mathrm{dF}}⸳FABP4$ | Degradation of FABP4 protein |

**Table S5.** Description of the model parameters and their values

| Description | Parameter | Values | Unit |
| --- | --- | --- | --- |
| Synthesis rate constant of Bmal1 ̶ Clock complex regulated through Per ̶ Cry pathway | $\boldsymbol{k}_{\boldsymbol{sbmal}\boldsymbol{1}\boldsymbol{p}}$ | **0.06** | **s.u. min^-1^** |
| Michaelis – Menten constant associated with Bmal1 ̶ Clock complex synthesis through Per ̶ Cry pathway | $\boldsymbol{J}_{\boldsymbol{bmal}\boldsymbol{1}\boldsymbol{p}}$ | **2** | **̶** |
| Per ̶ Cry complex mediated inhibition threshold | $\boldsymbol{k}_{\boldsymbol{mpercry}}$ | **0.007** | **s.u.** |
| Synthesis rate constant of Bmal1 ̶ Clock complex regulated through Rev pathway | $\boldsymbol{k}_{\boldsymbol{sbmal}\boldsymbol{1}\boldsymbol{r}}$ | **0.001** | **s.u. min^-1^** |
| Michaelis – Menten constant associated with Bmal1 ̶ Clock complex synthesis through Rev pathway | $\boldsymbol{J}_{\boldsymbol{bmal}\boldsymbol{1}\boldsymbol{r}}$ | **1** | **̶** |
| Rev mediated inhibition threshold | $\boldsymbol{k}_{\boldsymbol{mrev}}$ | **0.005** | **s.u.** |
| Degradation rate constant of Bmal1 ̶ Clock complex | $\boldsymbol{k}_{\boldsymbol{dbmal}\boldsymbol{1}}$ | **0.0031** | **min^-1^** |
| Basal transcription rate constant of Per mRNA | $\boldsymbol{b}_{\boldsymbol{mper}}$ | **0.1** | **s.u. min^-1^** |
| Bmal1 ̶ Clock complex mediated transcription rate constant of Per mRNA | $\boldsymbol{k}_{\boldsymbol{smper}}$ | **0.25** | **min^-1^** |
| GC mediated transcription rate constant of Per mRNA | $\boldsymbol{k}_{\boldsymbol{GC}}$ | **0.008** | **min^-1^** |
| Degradation rate constant of Per mRNA | $\boldsymbol{k}_{\boldsymbol{dmper}}$ | **0.25** | **min^-1^** |
| Translation rate constant of Per protein | $\boldsymbol{k}_{\boldsymbol{sper}}$ | **0.2156** | **min^-1^** |
| Association rate constant of Per ̶ Cry complex | $\boldsymbol{k}_{\boldsymbol{a}}$ | **0.1415** | **min^-1^** |
| Dissociation rate constant of Per ̶ Cry complex | $\boldsymbol{k}_{\boldsymbol{d}}$ | **0.004716** | **min^-1^** |
| Degradation rate constant of Per protein | $\boldsymbol{k}_{\boldsymbol{dper}}$ | **0.03** | **min^-1^** |
| Non-linear degradation rate constant of Per protein | $\boldsymbol{k}_{\boldsymbol{per}}$ | **3.2478** | **min^-1^** |
| Degradation rate constant of Per ̶ Cry complex | $\boldsymbol{k}_{\boldsymbol{dpercry}}$ | **0.015** | **min^-1^** |
| Non-linear degradation rate constant of Per ̶ Cry complex | $\boldsymbol{k}_{\boldsymbol{percry}}$ | **0.003773** | **min^-1^** |
| Degradation saturation constant | $\boldsymbol{J}$ | **0.04** | **̶** |
| Per dependent degradation weight | $\boldsymbol{q}$ | **1** | **s.u.^-1^** |
| Per ̶ Cry complex dependent degradation weight | $\boldsymbol{r}$ | **2.2** | **s.u.^-1^** |
| Basal synthesis rate constant of GC | $\boldsymbol{b}_{\boldsymbol{GC}}$ | **0.005** | **s.u. min^-1^** |
| Synthesis rate constant of GC regulated through Per pathway | $\boldsymbol{k}_{\boldsymbol{sGC}}$ | **0.015** | **s.u. min^-1^** |
| Hill constant associated with GC synthesis through Per pathway | $\boldsymbol{J}_{\boldsymbol{GC}}$ | **1** | **̶** |
| Per protein mediated inhibition threshold | $\boldsymbol{k}_{\boldsymbol{mGC}}$ | **0.05** | **s.u.** |
| Hill coefficient associated with GC synthesis through Per pathway | $\boldsymbol{m}$ | **2** | **̶** |
| Degradation rate constant of GC | $\boldsymbol{k}_{\boldsymbol{dGC}}$ | **0.01** | **min^-1^** |
| Basal transcription rate constant of Rev mRNA | $\boldsymbol{b}_{\boldsymbol{mrev}}$ | **0.1** | **s.u. min^-1^** |
| Bmal1 ̶ Clock complex mediated transcription rate constant of Rev mRNA | $\boldsymbol{k}_{\boldsymbol{smrev}}$ | **0.4** | **min^-1^** |
| GC mediated transcription rate constant of Rev mRNA | $\boldsymbol{k}_{\boldsymbol{rGC}}$ | **0.2** | **min^-1^** |
| Degradational rate constant of Rev mRNA | $\boldsymbol{k}_{\boldsymbol{dmrev}}$ | **0.25** | **min^-1^** |
| Translation rate constant of Rev protein | $\boldsymbol{k}_{\boldsymbol{srev}}$ | **0.1** | **s.u. min^-1^** |
| Hill constant associated with translation of Rev protein | $\boldsymbol{J}_{\boldsymbol{rev}}$ | **2** | **s.u.** |
| Hill coefficient associated with translation of Rev protein | $\boldsymbol{h}$ | **2** | **̶** |
| Degradation rate constant of Rev protein | $\boldsymbol{k}_{\boldsymbol{drev}}$ | **0.35** | **s.u. min^-1^** |
| Michaelis – Menten constant associated with degradation of Rev protein | $\boldsymbol{J}_{\boldsymbol{drev}}$ | **5** | **s.u.** |
| Basal transcription rate constant of PPAR$\boldsymbol{\gamma}$ mRNA | $\boldsymbol{k}_{\boldsymbol{sP}}$ | **0.619** | **s.u. min^-1^** |
| CEBPA mediated transcription rate constant of PPAR$\boldsymbol{\gamma}$ mRNA | $\boldsymbol{k}_{\boldsymbol{P}\boldsymbol{1}}$ | **0.99** | **s.u. min^-1^** |
| Hill constant associated with CEBPA mediated transcription of PPAR$\boldsymbol{\gamma}$ mRNA | $\boldsymbol{K}_{\boldsymbol{mP}\boldsymbol{1}}$ | **6.5** | **s.u.** |
| Hill coefficient associated with transcription of PPAR$\boldsymbol{\gamma}$ mRNA, CEBPA mRNA and FABP4 mRNA | $\boldsymbol{n}$ | **2** | **̶** |
| FABP4 mediated transcription rate constant of PPAR$\boldsymbol{\gamma}$ mRNA | $\boldsymbol{k}_{\boldsymbol{P}\boldsymbol{2}}$ | **1.21** | **s.u. min^-1^** |
| Hill constant associated with FABP4 mediated transcription of PPAR$\boldsymbol{\gamma}$ mRNA | $\boldsymbol{K}_{\boldsymbol{mP}\boldsymbol{2}}$ | **12.3** | **s.u.** |
| Degradation rate constant of PPAR$\boldsymbol{\gamma}$ mRNA | $\boldsymbol{k}_{\boldsymbol{dmP}}$ | **0.9** | **min^-1^** |
| Translation rate constant of PPAR$\boldsymbol{\gamma}$ protein | $\boldsymbol{k}_{\boldsymbol{tP}}$ | **0.5** | **min^-1^** |
| Basal activation rate constant of PPAR$\boldsymbol{\gamma}$ protein (stimulus-independent) | $\boldsymbol{k}_{\boldsymbol{0}}$ | **140** | **min^-1^** |
| Stimulus-induced activation rate constant of PPAR$\boldsymbol{\gamma}$ protein | $\boldsymbol{k}_{\boldsymbol{act}}$ | **20** | **min^-1^** |
| Stimulus-induced effective activation rate constant of PPAR$\boldsymbol{\gamma}$ protein | $\boldsymbol{k'}_{\boldsymbol{act}}$ | $\mathbf{k}_{\mathbf{act}}\mathbf{e}^{\left( \mathbf{k}\mathbf{T}_{\mathbf{p}} \right)^{\mathbf{n}_{\mathbf{p}}}}$ | **min^-1^** |
| Sensitivity parameter to pulse duration | $\boldsymbol{k}$ | **0.00054** | **min^-1^** |
| Degree of non-linearity for effective activation rate constant | $\boldsymbol{n}_{\boldsymbol{p}}$ | **1.4** | **̶** |
| Inactivation rate constant of PPAR$\boldsymbol{\gamma}$ protein | $\boldsymbol{k}_{\boldsymbol{inact}}$ | **4** | **min^-1^** |
| Degradation rate constant of PPAR$\boldsymbol{\gamma}$ protein | $\boldsymbol{k}_{\boldsymbol{dP}}$ | **0.7** | **min^-1^** |
| Basal transcription rate constant of CEBPA mRNA | $\boldsymbol{b}_{\boldsymbol{mCEBPA}}$ | **0.016** | **s.u. min^-1^** |
| PPAR$\boldsymbol{\gamma}$ mediated transcription rate constant of CEBPA mRNA | $\boldsymbol{k}_{\boldsymbol{C}}$ | **39.36** | **s.u. min^-1^** |
| Hill constant associated with PPAR$\boldsymbol{\gamma}$ mediated transcription of CEBPA mRNA | $\boldsymbol{K}_{\boldsymbol{mC}}$ | **10** | **s.u.** |
| Degradation rate constant of CEBPA mRNA | $\boldsymbol{k}_{\boldsymbol{dmC}}$ | **0.1** | **min^-1^** |
| Translation rate constant of CEBPA protein | $\boldsymbol{k}_{\boldsymbol{tC}}$ | **0.1** | **min^-1^** |
| Degradation rate constant of CEBPA protein | $\boldsymbol{k}_{\boldsymbol{dC}}$ | **0.07** | **min^-1^** |
| Basal transcription rate constant of FABP4 mRNA | $\boldsymbol{b}_{\boldsymbol{mFABP}\boldsymbol{4}}$ | **0.01** | **s.u. min^-1^** |
| PPAR$\boldsymbol{\gamma}$ mediated transcription rate constant of FABP4 mRNA | $\boldsymbol{k}_{\boldsymbol{F}}$ | **6.9** | **s.u. min^-1^** |
| Hill constant associated with PPAR$\boldsymbol{\gamma}$ mediated transcription of FABP4 mRNA | $\boldsymbol{K}_{\boldsymbol{mF}}$ | **20** | **s.u.** |
| Degradation rate constant of FABP4 mRNA | $\boldsymbol{k}_{\boldsymbol{dmF}}$ | **0.006** | **min^-1^** |
| Translation rate constant of FABP4 protein | $\boldsymbol{k}_{\boldsymbol{tF}}$ | **0.15** | **min^-1^** |
| Degradation rate constant of FABP4 protein | $\boldsymbol{k}_{\boldsymbol{dF}}$ | **0.11** | **min^-1^** |
| Glucocorticoid Stimulus | ***Stimulus*** | $\mathbf{4.0}\boldsymbol{\times}\frac{\frac{\mathbf{GC}}{\mathbf{sfcirc}}\boldsymbol{-}\mathbf{0.4}}{\mathbf{2-0.4}}$ | **̶** |
| Rate regulatory scaling factor of circadian network | $\boldsymbol{x}$ | **0.728** | **̶** |
| Rate regulatory scaling factor of adipogenic network | $\boldsymbol{y}$ | **1** | **̶** |
| Scaling factor of all proteins and mRNAs of circadian network | $\boldsymbol{sfcirc}$ | **1000** | **molecules** |
| Individual scaling factor of Per protein | $\boldsymbol{sfper}$ | **10** | **̶** |
| Individual scaling factor of Rev protein | $\boldsymbol{sfrev}$ | **1.5** | **̶** |
| Individual scaling factor of Rev mRNA | $\boldsymbol{sfrev}\boldsymbol{1}$ | **0.5** | **̶** |
| Scaling factor of all proteins and mRNAs of adipogenic network | $\boldsymbol{sf}$ | **1000** | **molecules** |
| Scaling factor of all mRNAs of adipogenic network | $\boldsymbol{sf}\boldsymbol{1}$ | **0.1** | **̶** |
| Individual scaling factor of PPAR$\boldsymbol{\gamma}$ protein and mRNA | $\boldsymbol{sfP}$ | **1.2** | **̶** |
| Individual scaling factor of PPAR$\boldsymbol{\gamma}$ mRNA | $\boldsymbol{sfP}\boldsymbol{1}$ | **1.2** | **̶** |
| Individual scaling factor of CEBPA protein and mRNA | $\boldsymbol{sfC}$ | **0.25** | **̶** |
| Individual scaling factor of CEBPA mRNA | $\boldsymbol{sfC}\boldsymbol{1}$ | **4** | **̶** |
| Individual scaling factor of FABP4 protein and mRNA | $\boldsymbol{sfF}$ | **0.2** | **̶** |
| Individual scaling factor of FABP4 mRNA  (1 s.u. is considered as 1000 molecules) | $\boldsymbol{sfF}\boldsymbol{1}$ | **3** | **̶** |

For the experiments involving Rosiglitazone as the activator of PPAR$\gamma$ protein, equation 1, 2 and 3 from **Table S1** (similarly equation 8, 9 and 10 from **Table S2**) are replaced by the following equations.

Table S6. Revised Ordinary Differential Equations for the Adipogenic network through Rosiglitazone activation in Fig. 1C

$$\frac{\boldsymbol{dmPPAR\gamma}}{\mathbf{dt}}\mathbf{=sf}\mathbf{⸳}\mathbf{sf1}\mathbf{⸳}\mathbf{sfP}\mathbf{⸳}\mathbf{sfP1}\mathbf{⸳}\mathbf{y}\mathbf{⸳}\left( {\mathbf{k}_{\mathbf{1}}\mathbf{⸳}\mathbf{k'}_{\mathbf{rosi}}\mathbf{+k}}_{\mathbf{sP}}\mathbf{+}\frac{\mathbf{k}_{\mathbf{P1}}\mathbf{⸳}\mathbf{CEBP}\mathbf{A}^{\mathbf{n}}}{\left( \mathbf{sf}\mathbf{⸳}\mathbf{sfC}\mathbf{⸳}\mathbf{K}_{\mathbf{mP1}} \right)^{\mathbf{n}}\mathbf{+CEBP}\mathbf{A}^{\mathbf{n}}}\mathbf{+}\frac{\mathbf{k}_{\mathbf{P2}}\mathbf{⸳}\mathbf{FABP}\mathbf{4}^{\mathbf{n}}}{\left( \mathbf{sf}\mathbf{⸳}\mathbf{sfF}\mathbf{⸳}\mathbf{K}_{\mathbf{mP2}} \right)^{\mathbf{n}}\mathbf{+FABP}\mathbf{4}^{\mathbf{n}}} \right)\mathbf{-y}\mathbf{⸳}\mathbf{k}_{\mathbf{dmP}}\mathbf{⸳}\boldsymbol{mPPAR\gamma}$$

$$\frac{\mathbf{dPPAR}\boldsymbol{\gamma}_{\mathbf{i}}}{\mathbf{dt}}\mathbf{=}\frac{\mathbf{y}\mathbf{⸳}\mathbf{k}_{\mathbf{tP}}\mathbf{⸳}\boldsymbol{mPPAR\gamma}}{\mathbf{sf1}\mathbf{⸳}\mathbf{sfP1}}\mathbf{+y}\mathbf{⸳}\mathbf{k}_{\mathbf{inact}}\mathbf{⸳}\mathbf{PPAR}\boldsymbol{\gamma}_{\mathbf{a}}\mathbf{-y}\mathbf{⸳}\left( \mathbf{k}_{\mathbf{0}}\mathbf{+}\mathbf{k'}_{\mathbf{rosi}}\mathbf{⸳}\mathbf{Rosiglitazone} \right)\mathbf{⸳}\mathbf{PPAR}\boldsymbol{\gamma}_{\mathbf{i}}\mathbf{-y}\mathbf{⸳}\mathbf{k}_{\mathbf{dP}}\mathbf{⸳}\mathbf{PPAR}\boldsymbol{\gamma}_{\mathbf{i}}$$

$$\frac{\mathbf{dPPAR}\boldsymbol{\gamma}_{\mathbf{a}}}{\mathbf{dt}}\mathbf{=y}\mathbf{⸳}\left( \mathbf{k}_{\mathbf{0}}\mathbf{+}\mathbf{k'}_{\mathbf{rosi}}\mathbf{⸳}\mathbf{Rosiglitazone} \right)\mathbf{⸳}\mathbf{PPAR}\boldsymbol{\gamma}_{\mathbf{i}}\mathbf{-y}\mathbf{⸳}\mathbf{k}_{\mathbf{inact}}\mathbf{⸳}\mathbf{PPAR}\boldsymbol{\gamma}_{\mathbf{a}}\mathbf{-y}\mathbf{⸳}\mathbf{k}_{\mathbf{dP}}\mathbf{⸳}\mathbf{PPAR}\boldsymbol{\gamma}_{\mathbf{a}}$$

| **Table S7.** Revised reaction governing the Adipogenic network through Rosiglitazone activation in Fig. 1C | | | |
| --- | --- | --- | --- |
| **PPAR**$\boldsymbol{\gamma}$ **mRNA** | | | |
| R44 | $\underset{\to}{Rosiglitazone}mPPAR\gamma$ | $\mathrm{sf}⸳sf1⸳\mathrm{sfP}⸳sfP1⸳y⸳k_{1}⸳{k'}_{\mathrm{rosi}}$ | Rosiglitazone-induced transcription of PPAR$\gamma$ mRNA |
| **Activated PPAR**$\boldsymbol{\gamma}$ **protein** | | | |
| R32 | $PPAR\gamma_{i}\underset{\to}{Rosiglitazone}PPAR\gamma_{a}$ | $y⸳{k'}_{rosi}⸳\mathrm{Rosiglitazone}⸳\mathrm{PPAR}\gamma_{i}$ | Rosiglitazone-induced activation of PPAR$\gamma$ protein |

**Table S8.** Description of the revised model parameters and their values

| Description | Parameter | Values | Unit |
| --- | --- | --- | --- |
| Transcriptional sensitivity factor of PPAR$\boldsymbol{\gamma}$ mRNA due to Rosiglitazone activation | $\boldsymbol{k}_{\boldsymbol{1}}$ | **0.00011** | **s.u.** |
| Rosiglitazone-induced activation rate constant of PPAR$\boldsymbol{\gamma}$ protein | $\boldsymbol{k}_{\boldsymbol{rosi}}$ | **20** | **min^-1^** |
| Rosiglitazone-induced effective activation rate constant of PPAR$\boldsymbol{\gamma}$ protein | $\boldsymbol{k'}_{\boldsymbol{rosi}}$ | $\mathbf{k}_{\mathbf{rosi}}\mathbf{e}^{\left( \mathbf{k}\mathbf{T}_{\mathbf{p}} \right)^{\mathbf{n}_{\mathbf{p}}}}$ | **min^-1^** |
| Sensitivity parameter to pulse duration | $\boldsymbol{k}$ | **0.008** | **min^-1^** |
| Degree of non-linearity for effective activation rate constant | $\boldsymbol{n}_{\boldsymbol{p}}$ | **0.15** | **̶** |

As a complementary analysis to the model predictions presented in the main manuscript, we examined whether modest reductions in the transcriptional activities of the adipogenic regulators could alter differentiation outcome only under continuous glucocorticoid (GC) signal. For this purpose, the basal transcription rate of CEBPA or FABP4 was reduced by 5%, while the corresponding PPARγ mediated transcription rate was reduced by 1%.

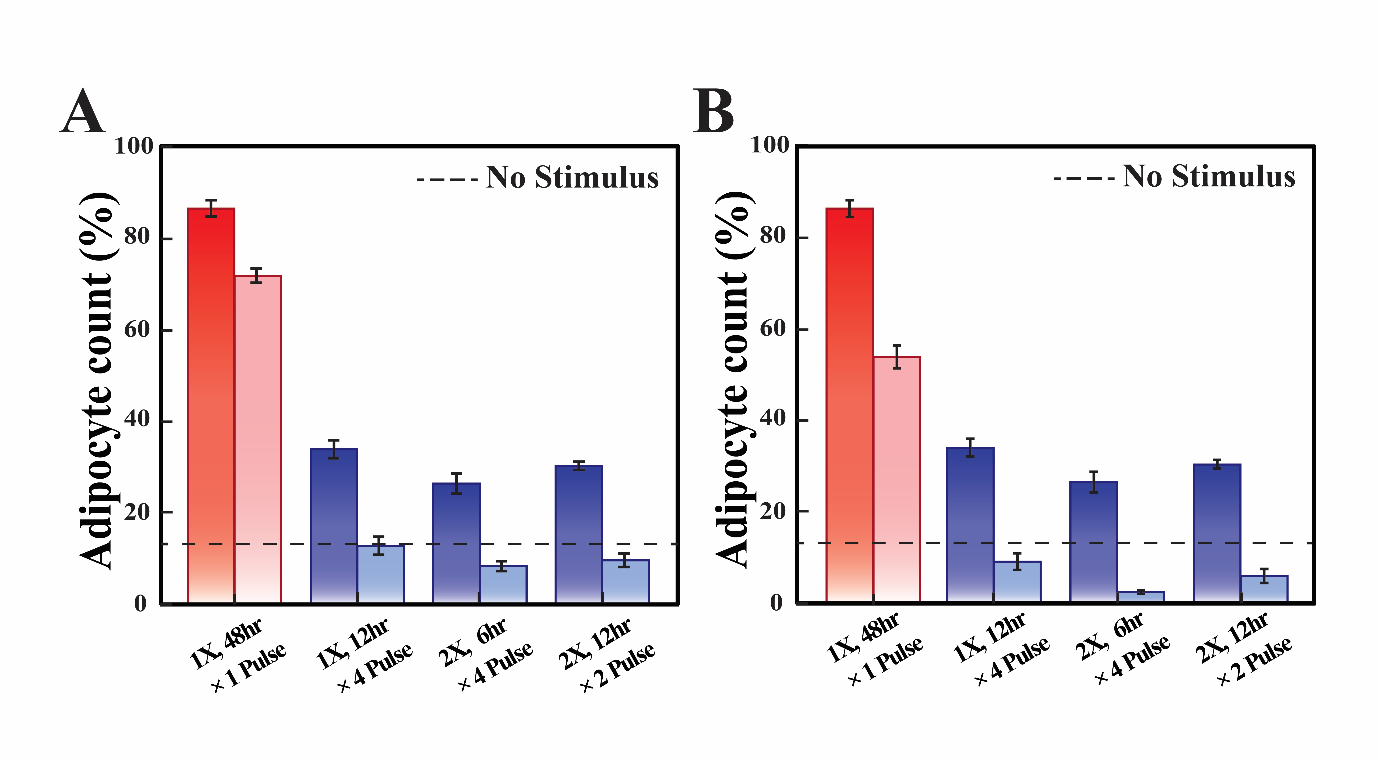

*Figure S1. Reduction in transcriptional activity attenuates adipogenic differentiation, while preserving the model’s ability to differentiate between continuous and pulsatile GC signals: (A) CEBPA, (B) FABP4.*

Reduced CEBPA transcriptional activity lowers adipocyte formation under all GC stimulation pattens (**Fig. S1**). A comparable reduction in differentiation is also observed when FABP4 transcriptional activity is decreased (**Fig. S1**). Nevertheless, continuous glucocorticoid stimulation remains considerably more effective at inducing adipogenesis than oscillatory glucocorticoid inputs. Thus, although reduced transcriptional activity attenuates adipogenic commitment, it does not abolish the model’s ability to distinguish between continuous and pulsatile GC signals. These findings suggest that suppressing individual components of the adipogenic program is insufficient to eliminate the temporal decoding of glucocorticoid dynamics.
